# Climbing fiber multi-innervation of mouse Purkinje dendrites with arborization common to human

**DOI:** 10.1101/2023.03.27.534425

**Authors:** Silas E. Busch, Christian Hansel

**Affiliations:** Department of Neurobiology, University of Chicago; Chicago, Illinois, 60637, USA

## Abstract

Canonically, each Purkinje cell in the adult cerebellum receives only one climbing fiber from the inferior olive. Underlying current theories of cerebellar function is the notion that this highly conserved one-to-one relationship renders Purkinje dendrites into a single computational compartment. However, we show that multiple primary dendrites are a near-universal morphological feature in human. Using tract-tracing, immunolabeling, and *in vitro* electrophysiology, we demonstrate in mice that ∼25% of mature polydendritic cells receive more than one climbing fiber input. Two-photon calcium imaging *in vivo* reveals that separate dendrites can exhibit distinct response properties to sensory stimulation, indicating some polydendritic cells integrate functionally independent climbing fiber receptive fields. These findings reveal that Purkinje cells are morphologically and functionally more diverse than previously thought.

---

Inputs to the cerebellar cortex are integrated by the dendrites of Purkinje cells (PCs), the sole cortical output neuron. Despite their well-characterized position in what is considered a conserved and stereotypical circuit (*1*), PCs exhibit remarkably diverse dendritic morphology in the rodent (*2*) and it is not known how specific features of dendritic arborization may affect their function.

Human PC morphology remains even more elusive. Studies of human PC morphology – dating back over 120 years to the iconic illustrations of Camillo Golgi and Santiago Ramón y Cajal (*3, 4*) – investigate small numbers of cells (*5–7*). While no information on frequency and distribution of morphological types is available, it can be observed that human PCs are often ‘polydendritic’, having either numerous trunks emerging from the soma or a proximal bifurcation of a single trunk. These features produce highly segregated dendritic compartments, raising the question whether this confers functional properties that have gone unreported.

Here, we specifically ask whether the existence of several primary dendrites enables multiple climbing fiber (CF) innervation in the adult cerebellum. During development, the early growth of a primary dendrite provides structural support for the ramification of a ‘winner’ CF amidst competitive elimination of surplus CFs (*8–10*). Weaker CF inputs fail to translocate to the dendrite, possibly as a result of competitive processes resembling adult bidirectional synaptic plasticity (*11–14*). In PCs where multiple primary dendrites conceivably offer a means to evade competition from other CFs, is the elimination pressure reduced enough to allow multiple CFs to be maintained? Would multi-innervation provide functionally independent receptive fields to distinct dendritic compartments? Here, we categorize PC morphology in human tissue and use a range of experimental methods in the mouse to characterize the relationship between PC morphology and CF input.

## A majority of human, but not murine, PCs have multiple primary dendrites

We used fluorescent calbindin immunolabeling to visualize PCs in post-mortem human tissue (Fig 1A). Based on proximal primary dendrite structure, which articulates the contours of the entire arbor, we define one standard structural category, *Normative* – one primary dendrite that may have a distant bifurcation (beyond a two somatic diameter threshold of 40µm in mouse and 50-70µm in human), and two polydendritic categories: *Split* – one trunk that bifurcates into multiple primary dendrites proximal to the soma (below the somatic diameter threshold); and *Poly* – multiple trunks emerging directly from the soma (Fig. 1A and fig. S1; see materials and methods). While these categories translate to the mouse (Fig. 1D), we find that mice diverge significantly from humans in that they have fewer Split PCs (36.6 vs. 47.7%) and far fewer Poly PCs (17.46 vs. 46.53%; Fig. 1G and fig S2A). Instead, in mice Normative PCs constitute the largest PC category (45.9%) in contrast to humans (5.8%).

**Fig. 1.**
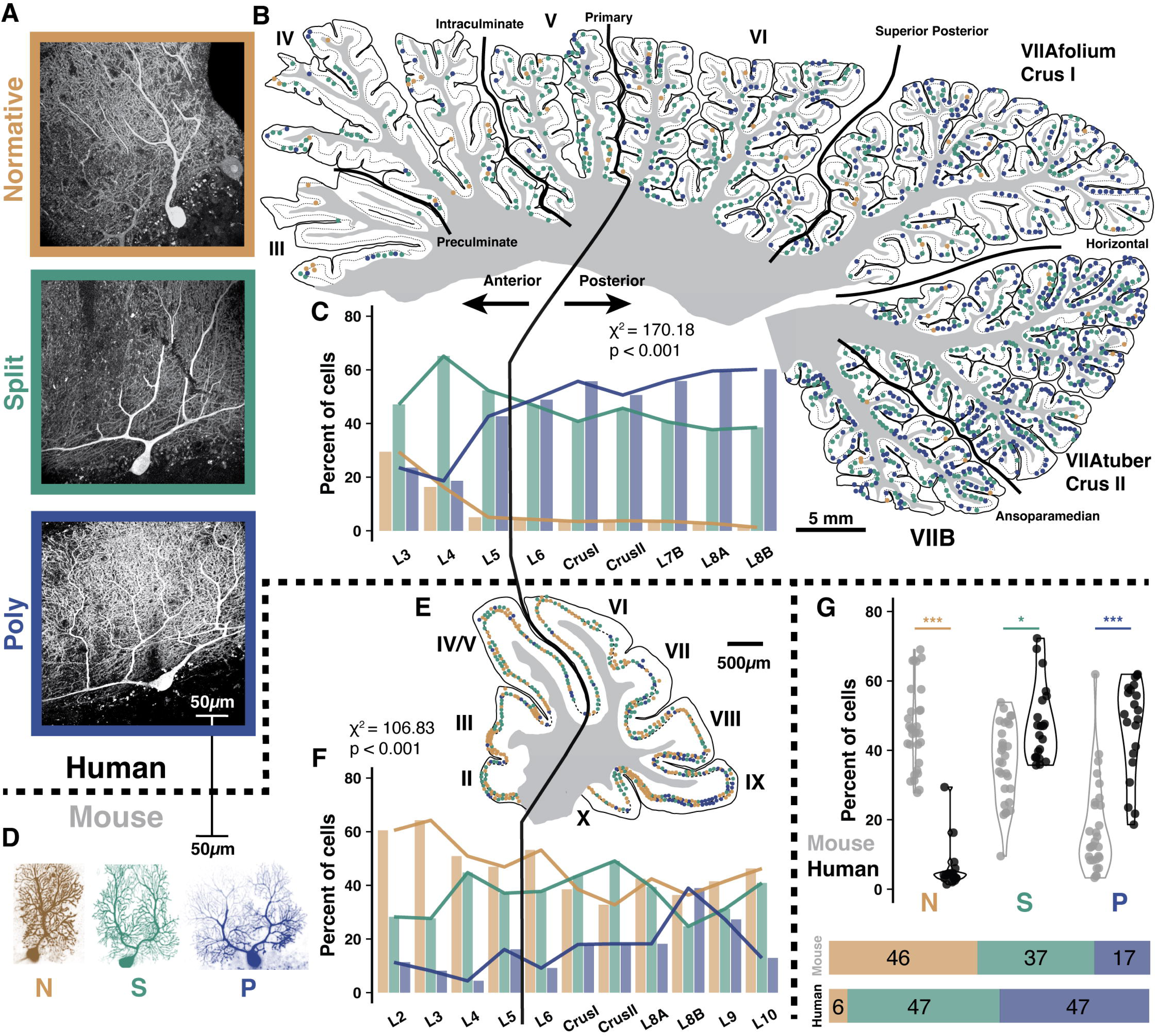
Comparative morphology and regional variability in human and mouse cerebellar Purkinje cells. (**A**) Immunolabeling of Purkinje cells in human reveals a range of dendritic morphologies, categorized by primary dendrite geometry as Normative, Split, or Poly. (**B**) Human mid-hemisphere reconstruction demonstrating the spatial distributions of each morphological type. Due to variable preservation of the tissue, some anterior lobules and intervening posterior sub-lobules had a lower density of labeled PCs. (**C**) Morphology demographics across lobules (n = 3 individuals >86yo, 6,640 cells; see table 1). (**D**) Purkinje cells filled with dye during a patch experiment in the mouse, to scale with human cells, also exhibit Normative, Split, and Poly morphology. (**E-F**) As in (**B-C**), but in mouse. (n = 3 mice >P50, 1,350 cells; see table 2). (**G**) Morphological category distribution counted by lobule in human (n = 20, 21, and 21 lobules) and mouse (n = 30, 30, 29) reveals a consistent increase in the number of Split and Poly PCs in human. *P < 0.05, **P < 0.01, ***P < 0.001.

We manually marked the distribution of dendritic morphologies in collectively ∼8,000 cells across whole, reconstructed parasagittal slices of mid-hemisphere in human and mouse (Fig. 1B,E and fig. S2B-C). In posterior lobules of human, there is a higher percentage of Poly PCs (52.04 vs. 29.86%) and a lower percentage of Normative PCs (3.52 vs. 12.71%) than in anterior lobules (Fig. 1C and fig S2A and table 1). While the total rate is far lower, Poly PCs are relatively more prevalent in posterior lobules of the mouse as well (10.61 vs. 21.65%, Fig. 1F-G and fig S2A and table 2).

The distribution observed does not appear to align with zonal or foliar patterning by zebrin or parvalbumin expression (*15*). Since the physiological implications cannot be readily studied in human, we turn to the corresponding mouse cells for further characterization.

## Multiple climbing fibers may innervate separate primary dendrites

CF activity causes complex spike firing in Purkinje cells (*16, 17*), which is reciprocally related to simple spike firing (*17, 18*) and exerts powerful control over dendritic integration and PF plasticity (*19–26*). Though many studies cite the critical importance of one-to-one CF to PC connectivity in cerebellar function – and abnormal connectivity in dysfunction – some work has shown CF multi-innervation in ∼15% of PCs in adult rodents (*27–29*). One study found elevated CF multi-innervation in sulcal regions of the folia, and suggested that dendrite structure may play a role (*29*). *Putative* multi-innervation has been linked to remodeling dendrite planarity in development (*30*), and is observed following genetic perturbations that leave inter-CF competition intact (*31, 32*).

To test whether multiple CF innervation can be found in mature PCs, we combined a sparse dextran tracer (DA-594) labeling of inferior olivary (IO) neurons (Fig. 2A) with immunolabeling of CF terminal boutons (VGluT2) and PCs (calbindin). As all CF terminals are marked by VGluT2, but only some will express DA-594, this method allows for the identification of multiple CF inputs from distinct IO neurons onto single Purkinje cells (*32*). Fig. 2B shows a Poly PC (P87) that is indeed innervated by two CFs on its separate primary dendrites.

**Fig. 2.**
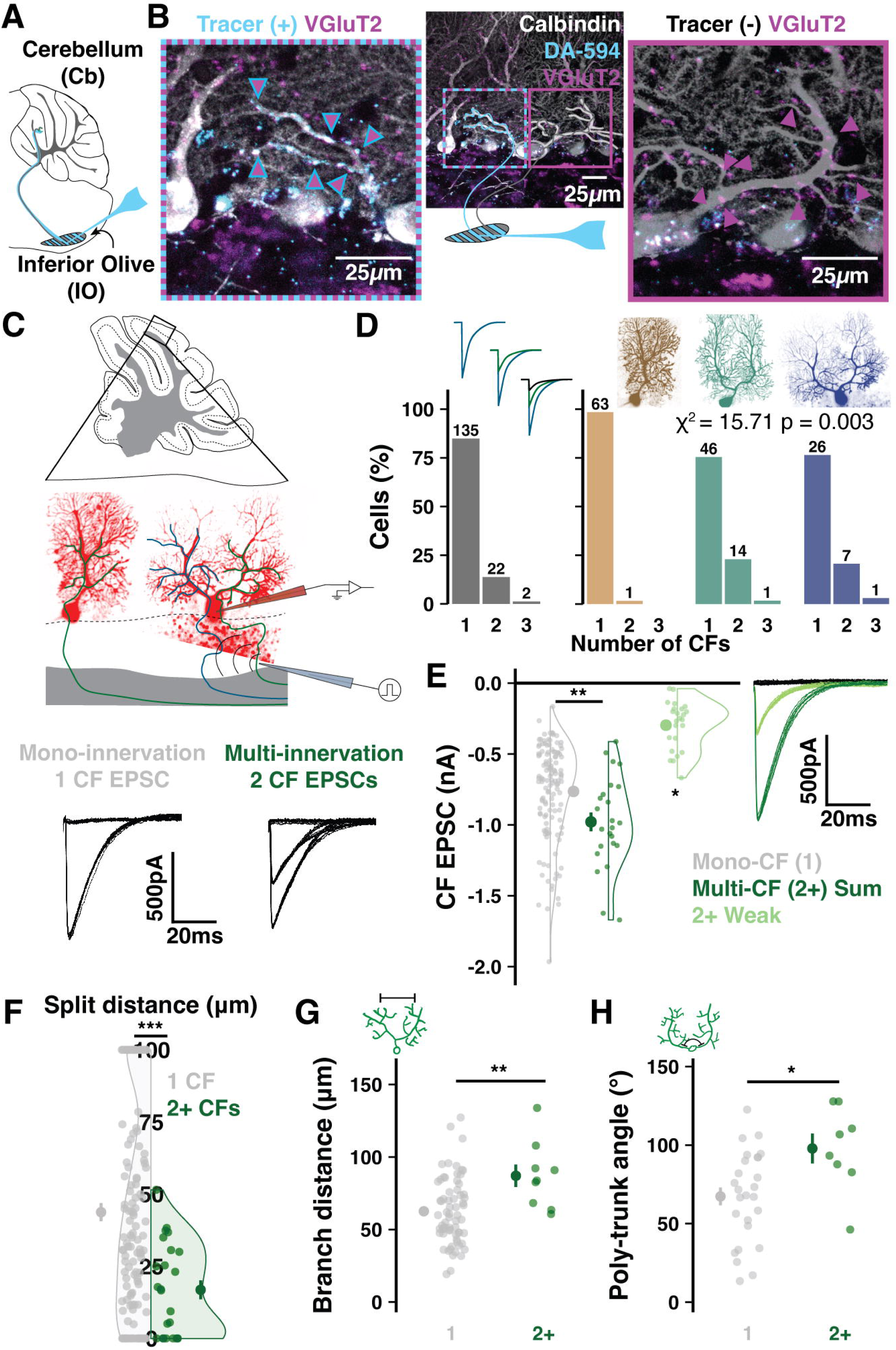
Climbing fiber multi-innervation of mature polydendritic Purkinje cells. (**A**) Schematic of tracer (DA-594) injection. (**B**) A Poly PC after immunolabeling for PCs (calbindin) and CF terminals (VGluT2). Tracer label distinguishes CFs with distinct olivary origin on the left and right trunks. (**C**) Scheme of whole-cell patch-clamp in cerebellar slices and CF EPSCs recorded from either a mono-or multi-innervated PC. (**D**) Number of mono-vs. multi-innervated PCs as a combined population (*left*). Categorizing by morphology reveals that effectively all multi-innervation occurs in polydendritic PCs (n = 50 animals, 159 cells). (**E**) Summed multi-CF EPSCs are larger than mono-CF EPSCs (n = 135 and 24 cells). The weaker of multiple CFs typically provides >200pA signals. Holding potential: −10 to −30mV (n = 24 cells). (**F**) Multi-innervated PCs have earlier dendrite bifurcations (n = 135 and 24 cells). (**G**) Among PCs with a bifurcated primary dendrite, multi-innervated cells have a wider distance between compartments (n = 85 and 9 cells). (**H**) Multi-innervated Poly PCs have a wider angle between emerging trunks (n = 26 and 8 cells). Summary points indicate mean ± SEM. *P < 0.05, **P < 0.01, ***P < 0.001.

## Quantification of CF multi-innervation in mature PCs

We obtained a quantitative measure of CF multi-innervation across the PC population by using whole-cell patch-clamp recordings in murine cerebellar slices (Fig. 2C). We adjusted current intensity and stimulus electrode position in the granule cell layer – sub-adjacent to each patched PC – to identify any ascending CF inputs and their stimulus thresholds. Mono-innervated PCs have a single, discrete EPSC while multi-innervated PCs exhibit two or more discrete EPSC amplitudes selectively evoked by distinct stimulus intensities (Fig. 2C bottom and fig. S3A-B). We observe that ∼15% of all PCs in mature animals (P20-66) receive multiple CFs (Fig. 2D). Within this cohort, we do not observe higher rates of multi-innervation in younger animals, confirming that CF competition for survival is complete by P20 (fig. S4L). CF multi-innervation is present across cerebellar regions and foliar sub-areas (fig. S4A-I). Posterior lobules have a higher frequency of multi-innervation (fig. S4H), possibly due to increased prevalence of Poly-PCs (fig. S4I-J), matching our finding in immunolabeled tissue (Fig. 1F-G).

Combining this technique with fluorescent dye loading and confocal imaging reveals that multi-innervation is largely restricted to PCs with polydendritic structures (23/24 PCs) and occurs in ∼25% of cells in this group (1/64 Normative, 15/61 Split, and 8/34 Poly PCs, Fig. 2D). The summed CF EPSC of multi-innervated PCs is larger, on average, than the amplitude of individual CF inputs to mono-innervated PCs. The amplitude of the smaller CF (at −30 to −10mV holding potential) is typically >200pA (Fig. 2E). This indicates that, under physiological membrane potentials, even the weakest of multiple CFs will likely deliver sufficient current to the soma to influence output (*33*). The amplitude of weaker CFs increased with age (fig. S3M), which may denote a delayed or elongated maturation period of these inputs relative to the completed development of single CF inputs or the more dominant of multiple CFs (fig. S3N).

The relative EPSC amplitude ratio between dominant and smaller CFs varies widely, but smaller CFs most often have >25% the relative amplitude of the dominant CF (Fig. S3O). This ratio differs across foliar sub-areas (fig. S3P) and correlates with the angle between Poly PC trunks (fig. S3Q), further emphasizing the relationship between morphology and CF input properties.

The prevalence of multi-innervation is correlated with proximity of bifurcation and angle of separation between emerging trunks in Split and Poly PCs, respectively (Fig. 2D-F and fig. S3D-F). Multi-CF PCs also have wider dendritic arbors in the parasagittal plane, while not differing in the angle of bifurcation (fig. S2D) or soma size (fig. S3H).

## CF multi-innervation produces heterogeneous Ca^2+^ signals across dendrites *in vivo*

Do multiple converging CFs provide functionally distinct input to a single Purkinje cell? How would this impact dendritic signaling *in vivo*? CF input triggers massive Ca^2+^ entry into PC dendrites via voltage-gated Ca^2+^ channels (*34*), NMDA receptors (*35*), and release from internal stores (*36*), which is subject to modulation by local ion conductances and their plasticity (*34, 37*) and local inhibition from interneurons (*38, 39*). These processes exhibit heterogeneity across branches (*40–42*), yet observations of branch-specific, CF-dependent Ca^2+^ signals have not been reported *in vitro* and are rare *in vivo* (*41, 42*). Branch-specific Ca^2+^ signaling would provide a mechanism for large compartments to conduct independent integration of input.

We obtained a sparse PC expression of the Ca^2+^ indicator GCaMP6f and used two-photon imaging of awake mice to record non-evoked ‘spontaneous’ Ca^2+^ signals from primary dendrite compartments in small populations of <10 cells (Fig. 3A-B; see materials and methods). In this configuration, non-evoked Ca^2+^ signals beyond the micro-compartment scale are almost entirely CF-dependent (*43, 44*) and Ca^2+^ event amplitude reflects the number of spikes in the presynaptic CF burst (*45*). This is confirmed by our observed ∼1.2Hz spontaneous Ca^2+^ event frequency (fig. S5D) that matches an expected CF input frequency moderately greater than 1Hz (*17*). Fluorescence traces for each primary dendrite were extracted and deconvolved separately to contrast event amplitude and frequency across branches (Fig. 3C-E).

**Fig. 3.**
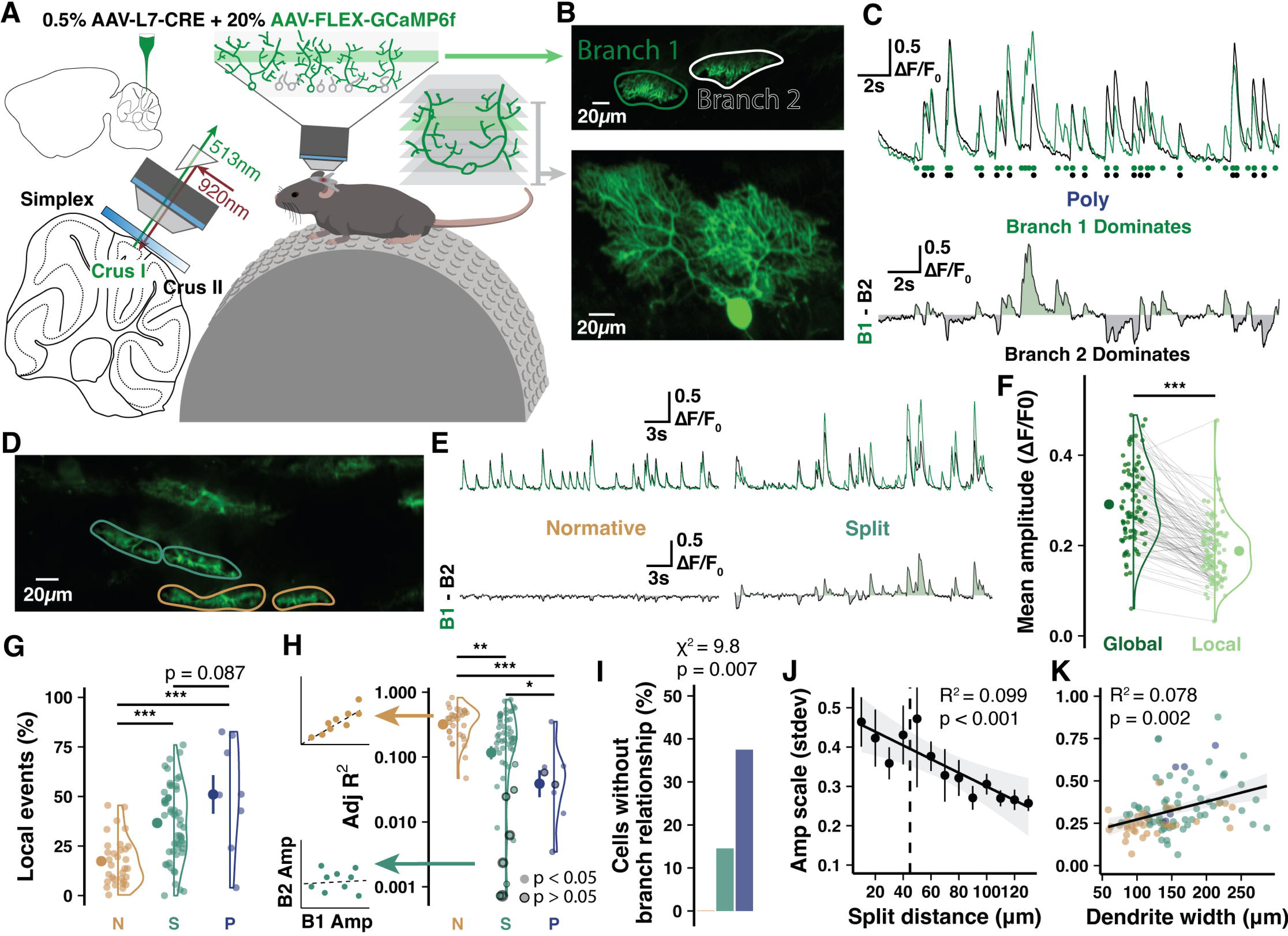
Two-photon imaging *in vivo* reveals Ca^2+^ signal heterogeneity across Purkinje cell dendrites. (**A**) Schematic of experimental preparation. (**B**) Example imaging plane and 3D-reconstruction of a Poly PC. (**C**) Spontaneous signal and deconvolved events (circles) by branch with difference trace below demonstrates heterogeneous global event amplitude scale and branch-specific events. (**D-F**) Another recording from a Normative and Split PC highlights homogeneous vs. heterogeneous signaling. (**F**) Local events are moderately smaller than global events (n = 15 animals, 95 cells). (**G**) Branch-specific local events as a percentage of total events in each cell by morphology (n = 15 animals; n = 32, 55, and 8 cells). (**H**) Linear regressions on branch cross-correlation quantifies branch similarity (*left*). Model fit R^2^ values (*right*) reveals that cells with low branch signal similarity are predominantly Split and Poly PCs (n = 32, 55, and 8 cells). Bordered points indicate non-significant covariance. (**I**) Cells lacking detectable relationship using regression on inter-branch amplitudes are all Split or Poly PCs. (**J-K**) Interbranch amplitude variation by split distance (n = 105 cells) and total parasagittal width of the dendrites (n = 109 cells). Summary points indicate mean ± SEM. *P < 0.05, **P < 0.01, ***P < 0.001.

In each cell, we first identified local Ca^2+^ peaks detected in only one branch (Fig. 3C, F-G and Movie 1), which were moderately smaller than globally expressed events (Fig. 3F). We also compared the inter-event cross-correlation of global Ca^2+^ events expressed across branches, for which the fit and significance of a linear regression describes the inter-branch covariation (Fig. 3H-I).

As expected, most PCs have homogenous Ca^2+^ signals with linear inter-event covariance relationships across branches (Adj R^2^ > 0.1) and low numbers of local events (Fig. 3G-H). However, some PCs exhibited Ca^2+^ signal heterogeneity characterized by a linear regression of inter-event covariation with low Adj R^2^ < 0.1 that was not significant (0, 15, and 38% of Normative, Split, and Poly PCs, respectively; Fig. 3H-I) or a higher ratio of local events (17.4, 36.6, and 51%; Fig. 3G). High variability of inter-event amplitude scale between branches, another measure of heterogeneity, correlated with the bifurcation distance and total parasagittal dendritic width (Fig. 3J-K). This further links heterogeneity to underlying morphological contours defined by primary dendrite geometry.

Confirming that local events are the product of additional CF input, PCs with high local event rates had higher mean (fig. S5G-P) and maximum total event rates (fig. S5L-P), producing a larger dynamic range (fig. S5Q-R).

## Climbing fibers convey distinct whisker receptive fields to separate primary dendrites

To identify CF receptive fields (RFs) and their localization on PC dendrites, we took advantage of the discrete organization of whiskers as a sensory input array (*46, 47*). We anaesthetized animals to stimulate individual whiskers at 2Hz for 50s periods while recording Ca^2+^ activity of PCs in medial Crus I (Fig. 4A and S6A; see materials and methods). As expected, most PCs have only global events identically represented across primary dendrites. However, some PCs have high numbers of local events in response windows during the stimulus period (Fig. 4C) that can vary in magnitude between distinct whisker stimuli (Fig. 4D), indicating RF selectivity.

**Fig. 4.**
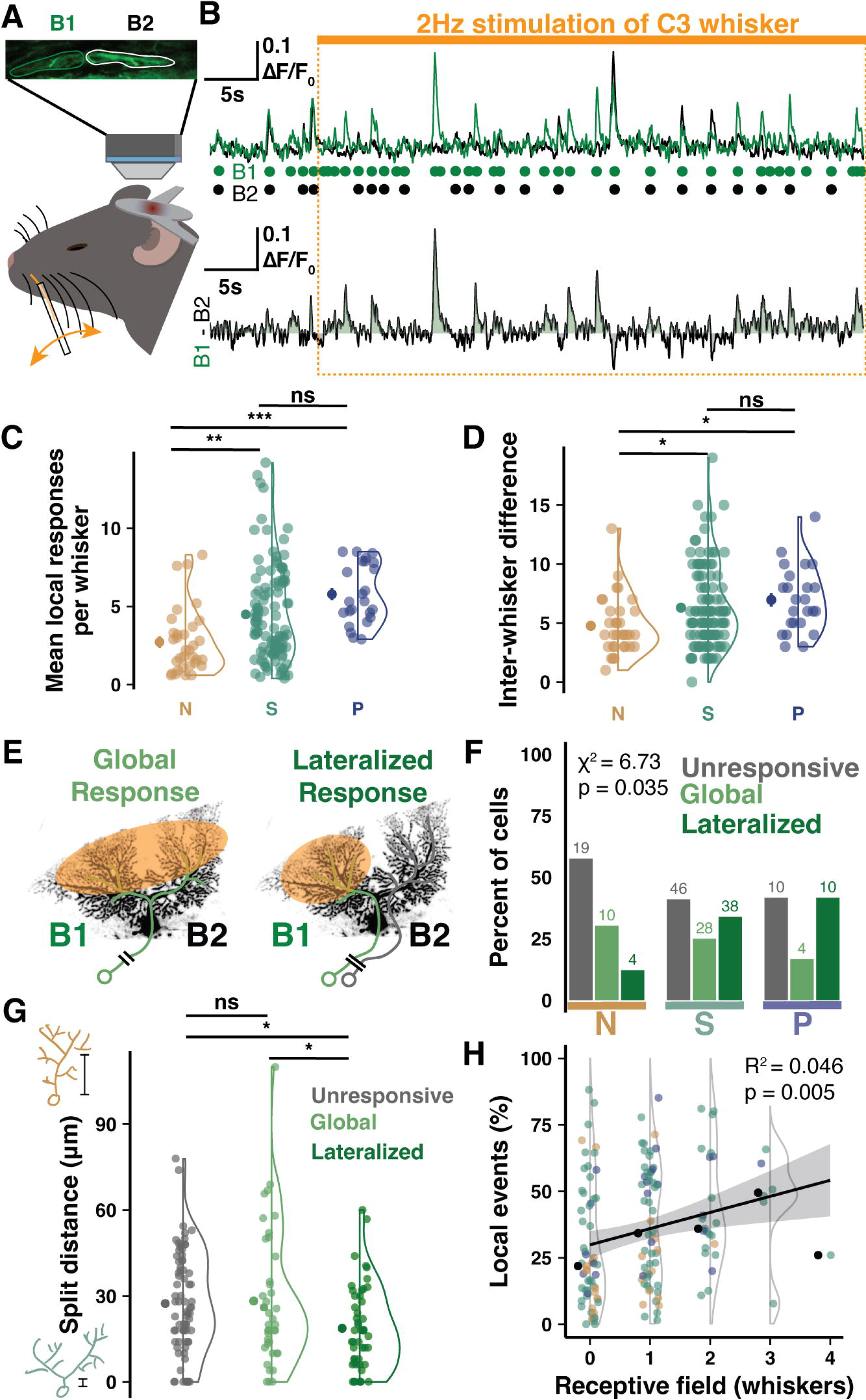
Branch-specific whisker receptive fields produced by climbing fiber multi-innervation of polydendritic Purkinje cells. (**A**) Schematic of the imaging configuration and whisker stimulation under anesthesia. (**B**) Sample traces and deconvolved events by branch during 50s whisker stimulation. Each whisker is tested twice; data from both periods are combined. Responsiveness of one branch and not the other drives an enhanced local event rate in B1 during the stimulus period. (**C**) Mean number of local branch events in response windows during stimulus periods of each tested whisker (n = 13 animals P95-120, n = 33, 112, and 24 cells). (**D**) Difference in local event number between whiskers eliciting maximum and minimum local responses (n = 33, 112, and 24 cells). (**E**) Schematic of global vs. lateralized responses. (**F**) Percentage of PCs by dendritic response profile and morphological category. Fewer Normative PCs have lateralized responses than polydendritic PCs (n = 169 cells). (**G**) Cells with lateralized responses have shorter split distances (n = 75, 42, and 52 cells). (**H**) Cells with more spontaneous local events respond to a higher number of whiskers (n = 151 cells). Summary points indicate mean ± SEM. *P < 0.05, **P < 0.01, ***P < 0.001.

Anaesthetized activity is sparsened, so responses were determined using the z-scored response probability during the experimentally bootstrapped high-frequency stimulus (fig. S6B-D). Using the z-scored response probability of each dendrite, we could observe a ‘lateralized’ response in some PCs, in which local events of one branch constitute a whisker response that is not observed in the other branch (Fig. 4E-F). Nearly all lateralized responses arise in Split and Poly PCs (48/52 cells, 92%; Fig. 4F) and in PCs with more signal heterogeneity, which map to Split and Poly PCs as in previous experiments (fig. S6F). Furthermore, PCs with lateralized responses have bifurcations that occur more proximally than PCs with only global responses (18.73 µm vs. 28.24 µm; Fig. 4G). Importantly, PCs with a higher ratio of branch-specific spontaneous events also exhibit responses to more whiskers, denoting an integration of more whiskers into their RFs (Fig. 4H). This supports the hypothesis that heterogeneous signals represent distinct, converging RFs such that heterogenous PCs are sampling more upstream RFs carried by functionally independent CF inputs.

## Climbing fiber induced branch-specific representations of stimulus modality in awake mice

While anesthesia provided excellent control and precision for single whisker stimulation, even subanesthetic ketamine alters network activity (*48*). To confirm that PC primary dendrites can differentially represent CF RFs in a more naturalistic state, we exposed awake animals to uni-and multisensory stimuli (Fig. 5A). As a major hub for sensory integration during associative learning, PC dendrites are an important model for how converging input profiles are represented across dendrites. The amplitude and duration of CF-induced dendritic Ca^2+^ spikes depend on stimulus strength (*44, 49*), which is reflected in CF burst behavior (*45*), and also on synaptic connectivity and weight of the CF input itself (*50*).

**Fig. 5.**
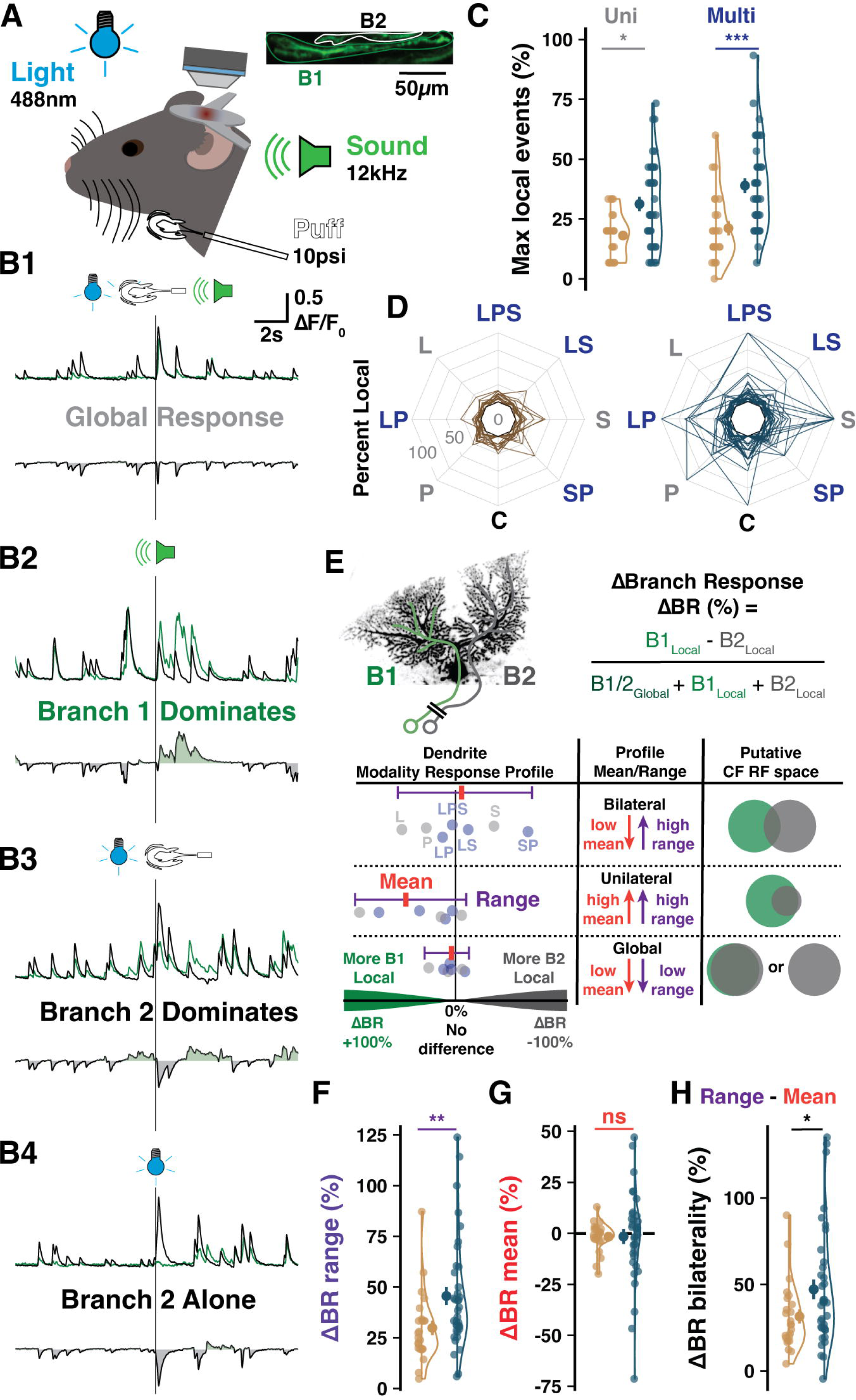
Branch-specific multisensory receptive fields. (**A**) Scheme of imaging and sensory stimulation of awake animals. (**B**) Sample traces showing combinations of inter-branch responses to different stimulus modalities. (**C**) The maximum number of local events observed for a stimulus of any category (here and below: n = 12 animals, n = 24 and 38 cells). (**D**) The percentage of responses having a local component, regardless of branch identity, across control (C), uni-, or multisensory trials in Normative vs. Split and Poly (S/P) PCs. Lines connect values for each PC. (**E**) Calculation of ΔBranch Response (ΔBR, *top*) between the stimulus types most favoring opposite branches. ΔBR values of each modality are calculated for each cell (*bottom, schematic points*) to map the ΔBR profile across stimuli and identify the mean and range. (**F**) The range is more pronounced in S/P cells (n = 24, 38). (**G**) No group difference in ΔBR mean (n = 24, 38). (**H**) Response profile bilaterality is the subtraction of ΔBR mean from the range. S/P PCs exhibit more bilaterality due to high ranges and low means (n = 24, 38). Summary points indicate mean ± SEM. *P < 0.05, **P < 0.01, ***P < 0.001.

We stimulated awake animals with light (488nm, ipsilateral), sound (12kHz tone, bilateral), and peri-oral air puff (10psi, ipsilateral) stimuli either alone or in multi-modal combinations while recording response properties in PC primary dendrites. Sensory evoked events – more than spontaneous – typically produce a global dendritic signal with consistent inter-trial amplitude ratio between branches (Fig. 5B1). Yet, we also observed complex sensory-evoked bursts of CF input with heterogeneous amplitudes between branches (Fig. 5B2-3) and either branch-specific responses alone (Fig. 5B4) or combined with a global response (Fig. 5B2). While PCs with multiple primary dendrites (Split and Poly, or S/P) have similar total response probabilities as Normative PCs (Fig. S7C), a larger share of responses were branch-specific in S/P PCs across stimulus modalities (Fig. 5C-D and S7A-B). To assess the relationship between uni- and multi-modal stimuli, we identified the maximum branch-specific responses to stimuli of each category (Fig. S7D-E), obtained the difference between uni- and multi-sensory maxima, and found an enhanced rate of local responses in S/P but not Normative PCs (Fig. S7F). This revealed that – as a more multifaceted change in the external world – multisensory stimuli could enhance the differential representation of CF RFs across primary dendrites in putatively multi-innervated PCs while failing to have an effect on mono-innervated Normative PCs.

While the previous analyses were blind to branch identity, we next asked how much the differential representation of each stimulus could favor one branch over the other. To do this, we generated a ΔBranch Response (ΔBR) index for each stimulus modality by calculating the difference in branch-specific, local responses as a fraction of total responses (Fig. 5E, *top*).

Absolute ΔBR indicates the reliability of local responses on either branch while the sign of the ΔBR indicates which branch over-represented the modality. This allowed us to generate a profile of branch-specific representation across all stimulus modalities, which could be quantified by the ΔBR mean and range for each cell (Fig. 5E, *bottom*). In this way, PCs could be distinguished as having one of three classes of multisensory response profile: *global*, with identical representation across branches in all cases; *unilateral*, with one branch exhibiting a larger RF representation than the other; and *bilateral*, with both branches capable of differentially representing unique stimulus modalities.

On average, S/P cells had a wider range, denoting branch-specific (e.g., unilateral or bilateral) representations that are more distinct across modalities (Fig. 5F and S7G-I). Cells for which only one branch exhibits local responses – unilateral – would have both a large ΔBR range but also a ΔBR mean that deviates from zero to favor that branch. To better characterize whether some PCs had bilateral representation profiles, we calculated the bilaterality of the RF profile by subtracting the ΔBR mean from the range. This showed that the local responses of S/P cells, more than Normative, produced RF profiles wherein a larger percentage of local signaling produced bilateral representations across sensory modalities (Fig. 5H and S7G-K). Collectively, this shows that PCs with multiple primary dendrites can differentially represent RFs of distinct CF inputs across their separate dendrites in the awake, mature mouse.

## Discussion

We show that non-canonical climbing fiber multi-innervation of Purkinje cells does occur in the mature murine cerebellum and is dependent on primary dendrite morphology. Nearly all observed multi-innervation occurs in neurons with multiple primary dendrites. Based on a quantitative categorization of >6,000 PCs from three human brains, we report that this type of PC dendritic structure is predominant in the human cerebellum. In contrast, we find that only a minority of murine PCs fall into the Split or Poly category. Within these morphological groups, about 25% of PCs are innervated by two or more CFs in the mouse.

Our two-photon recordings suggest that the majority of multi-innervated Purkinje cells have the capacity for branch-specific CF signaling and have distinct CF RFs. Our data do not allow us to conclude that the same results would be found in human PCs if such recordings were possible. However, they do describe a new motif in Purkinje cell dendritic compartmentalization: separate dendritic subfields with their own assigned CF inputs may emerge when early branching forms a polydendritic architecture.

CFs provide instructive signals in cerebellar function and plasticity (*17*) by encoding signals related to error (*19, 51*), sensory omission (*52*), as well as reward or reward-prediction (*53, 54*). Our findings constitute a substantial shift from the currently held belief that one CF innervates each PC. Instead, our observations suggest that one CF innervates each primary dendrite.

The consequences are potentially multifold, but one immediately results from geometric considerations. Multiple primary dendrites and their minor branches often fan out horizontally in the sagittal plane, forming a cleft between compartments. This configuration inevitably leads to a wider physical gap – and potential functional separation regarding encoded information content – between innervating PF bundles. Having a single cell body receive input from PF populations on multiple segregated dendrite structures may require – at least in a sub-population of these cells – innervation by distinct CFs to enable supervised learning based on matching RFs. Multiple CF innervation in the adult cerebellum thus preserves Purkinje cell function as a perceptron (*55*) with the increase of dendritic complexity. Upon somatic convergence, these two or more dendritic signals could assume different logic gate functions (e.g. AND, OR, XOR), depending on activation demands of target cells in the cerebellar nuclei (*56*).

## Materials and Methods

### Subjects

Human cerebellar tissue was collected from three embalmed donor bodies provided to the University of Chicago Pritzker School of Medicine Anatomy Lab by the Anatomical Gift Association of Illinois (AGAI). Individuals were 92 (F), 95 (F), and 86 (M) years old, died of causes unrelated to cerebellar morphology (e.g. ‘failure to thrive’, likely ‘failure to thrive’, and colon cancer, respectively), and tissue was stored for 2, 6, and 2 months, respectively. During life, all study subjects signed an informed consent approved by the AGAI.

For experiments involving mice, both *in vitro* and *in vivo*, all experimental and surgical procedures were in accordance with the University of Chicago Animal Care and Use Committee guidelines. We used wildtype C57BL/6J mice housed on a 12hr light/dark cycle. Animals of either sex were used in all experiments and no sex differences were observed in any reported measures.

### Immunohistochemistry

#### Embalmed human tissue

Due to incomplete fixation during the embalming process, we immediately submerged whole cerebella in 4% paraformaldehyde (PFA) for one week after they were obtained. Following this fixation period, each specimen was sectioned by hand in the sagittal axis to obtain 2-5mm blocks from the mid-hemisphere of each individual. Given the anterior curvature of folia in the hemisphere, blocks were cut at varying angles relative to the midline. Tissue blocks with incomplete fixation of deep structures were further fixed for 2-4 days. Depending on their size, blocks were cut transversely into dorsal and ventral sections, typically through the horizontal fissure, such that Lobules III through VIIAf/Crus I were in the dorsal block and VIIAt/Crus II through VIIIB in the ventral block. Occasionally, we also needed to cut the dorsal block into rostral and caudal sections. Next, each block was rinsed in 0.01M Phosphate buffer saline (PBS), dried on one side, mounted with super glue into the slicing chamber of a vibratome (Leica VT-1000S), and sliced at 30µm in the parasagittal plane.

Slices selected for immunolabeling were transferred to a clear tray, placed over a broad-spectrum LED array, covered with a reflective aluminum foil lid, and photobleached at 4L for 3-4 days. This reduced the strong autofluorescence in the green channel. Then tissue was washed in 50mM Glycine in 0.01M PBS for 2hrs at 4L and incubated in 20mM Sodium Citrate in 0.01M PBS at 50-60L using a heated water bath for 30min. After cooling to room temperature (RT), tissue was washed in 20mM Sodium Citrate for 5min then rinsed 2×30sec in dH20. Next, slices were permeabilized at RT in 0.01M PBS containing 0.025% Triton-X (PBS-TX) for 1hr. Blocking was done with PBS-TX containing 5% normal donkey serum (NDS) and 5% bovine serum albumin (BSA) for 1hr at RT followed by incubation in guinea pig anti-calbindin primary antibody (1:1000, Synaptic Systems) solution overnight (18-20hrs) at 4L with 1% normal donkey serum in PBS-TX. After 3×10min washes in PBS-TX at RT, slices were incubated in donkey anti-guinea pig AF488 secondary antibody (1:200, Jackson ImmunoResearch) for 2hrs at 4L with 1% NDS in PBS-TX. Finally, slices were washed in PBS-TX for 3×10min, mounted and coverslipped with Vectashield (Vector Laboratories, Inc.), and allowed to set overnight before visualization.

Slides were visualized under 10x or 20x magnification (Zeiss Achroplan 0.25NA, air; Olympus UMPlanFL N 0.5NA, water) and illuminated with an epi-fluorescent light source (LEJ HBO-100) cast through a 450nm-pass filter cube. This allowed us to manually scan through the cerebellar cortex and classify Purkinje cells (PCs) by their dendritic morphology. Post-mortem storage, embalming, and subsequent short-term submersion in ethanol, which renders many antigen sites inaccessible, diminished the tissue quality; however, this provided some advantage for our purposes. The condition of the tissue resulted in a sparse labeling of PCs that we expect to be random and without morphological bias. The sparsity provided a clearer visualization of each cell’s individual morphology and decreased the total number of cells so that an exhaustive count for each region was feasible and unbiased without the use of stereology.

To mark the morphology and cell location accurately, we initially traced the outlines of the pial surface, white matter tracts, and PC layer over low resolution images of the entire section. Cells were only included for categorization if the soma and at least 200µm lengths of primary dendritic trunks were clearly labeled such that all features of Normative, Split, and Poly categories were unambiguously present or absent (Fig. 1A and fig. S1). We marked the location and morphological type of each cell in the slice map and scanned these notes as an input image to a custom Matlab GUI. This allowed us to generate a .csv table output with a cell ID, XY coordinates, morphological category, region of the folia (gyrus, bank, or sulcus), and lobule of each marked cell (Fig. 1B and fig. S2). These data were imported to R for downstream analysis and plotting.

#### Purkinje cell morphological category definitions and criteria

In human, PCs were deemed Normative if they had the following features: 1) a single trunk emerging from the soma, and 2) either no bifurcation of the primary trunk within two soma distances (2x the diameter of the soma, 25-35µm per soma) or a highly asymmetrical bifurcation where the smaller branch did not project in the parasagittal axis more than 200µm from the main dendritic compartment. PCs were defined as Split if they had the following features: 1) a single trunk emerging from the soma, and 2) either symmetrical bifurcation of the primary trunk within two soma distances or an asymmetrical bifurcation within two soma distances where the smaller branch projected more than 200µm from the main dendritic compartment and thus reached prominence by its overall length and sub-branching. PCs were defined as Poly if they had more than one trunk emerging from the soma regardless of relative size.

In mouse, PC categories were defined the same way, except that the bifurcation threshold of two soma distances (each soma diameter is 18-22µm) was set at 40µm, and the smaller branch of an asymmetrical bifurcation had to project only 100µm away from the main dendritic compartment.

#### Climbing fiber tracer immunohistochemistry

Tracer injections in the inferior olive (Fig. 2A) were performed on mice aged 13-15 weeks under ketamine/xylazine anesthesia (100 and 10mg/kg, respectively, 0.1mL/10g weight, Covetrus) with subcutaneous injections of meloxicam (0.06mL, 1-2 mg/kg), buprenorphine (0.05mL, 0.1 mg/kg, Covetrus), and sterile saline (0.5-1mL). Body temperature was maintained at 35-37L with a feedback dependent heating pad. Mice were positioned in an upright, sitting position with the head clamped such that the line described by the maxilla and the ear bars in the acoustic foramen was parallel with ground. In this configuration, the atlanto-occipital joint membrane was exposed when skin above the posterior skull and posterior muscles attaching to the occipital bone were removed. Cutting open the membrane, approximately 1uL of 25% Alexa 594 conjugated dextran amine tracer (10,000 MW, Invitrogen) in saline was injected 1.8mm deep and 0.5mm lateral from midline into the medulla at a 59° angle (25). After 4-5 days of recovery, mice were anesthetized with ketamine/xylazine (100 and 10mg/kg) and perfused with 4% PFA. Cerebella were removed and incubated for 2hrs in 4% PFA at 4L and then overnight in 30% sucrose in 0.1M PB at 4L (until the tissue sank from the surface). The tissue was then rinsed briefly in 0.1M PB, dried and blocked, submerged in OCT medium, flash frozen, and then sliced (50μm, parasagittal plane) using a cryostat microtome (CM 3050S, Leica).

After slicing, tissue was immunolabeled as described above with several changes: glycine incubation for 1hr instead of 2hrs and heated Sodium Citrate incubation for 20min instead of 30min. Slices were incubated in primary antibody solution with rabbit anti-VGluT2 (1:500, Invitrogen) and guinea pig anti-calbindin (1:1000, Synaptic Systems), then in secondary antibodies with donkey anti-rabbit AF647 and donkey anti-guinea pig AF488 (both 1:200, Jackson ImmunoResearch). Slices were imaged at 40x (Zeiss EC Plan-Neofluar 1.3NA, oil immersion) and z-stacks of the molecular layer were obtained with a confocal microscope (Fig. 2B; Zeiss LSM 5 Exciter, Axioskop 2).

### Slice Electrophysiology

To quantify the frequency of functional climbing fiber (CF) multi-innervation, we used whole cell patch clamp and electrical stimulation in acute cerebellar slices with a cesium internal solution. Mice (P20-65) were anesthetized with isoflurane and decapitated. The cerebellum was immediately dissected in ice cold artificial cerebrospinal fluid (ACSF) containing (in mM): 124 NaCl, 5 KCl, 1.25 Na_2_HPO_4_, 2 CaCl_2_, 2 MgSO_4_, 26 NaHCO_3_, and 10 D-glucose, bubbled with 95% O_2_ and 5% CO_2_. Sagittal slices of the cerebellum (250μm thick, including medial hemisphere, paravermis, and vermis) were prepared with ice cold ACSF in a chilled slicing chamber using a vibratome (Leica VT-1000S), and allowed to recover for 1hr at room temperature in oxygenated ACSF. During recordings, the slices were continuously perfused with oxygenated ACSF containing 100μM picrotoxin to block GABA_A_ receptors.

Whole cell patch-clamp recordings from the PC somata were performed at room temperature using an EPC-10 amplifier (Fig. 2C and S3A; HEKA Electronics). The workstation was also equipped with a confocal microscope (Fig. 2C; LSM 5 Exciter and Axioskop 2, Zeiss) for the identification and morphological characterization of patched and dye-filled cells. Currents were filtered at 3kHz, digitized at 25kHz, and acquired using Patchmaster software (HEKA Electronics). For recordings of CF-EPSCs (Fig. 2C and S3A-B), the pipette solution was Cesium based to improve space clamp of inputs to distal dendrites and contained (in mM): 60 CsCl, 10 Cs D-Gluconate, 20 TEA-Cl, 20 BAPTA, 4 MgCl_2_, 4 Na_2_ATP, 0.4 Na_3_GTP, 30 HEPES (osmolarity: 295-305mmol/kg; pH 7.3, adjusted with CsOH). Alexa-633 dye (30μM) was added to the pipette solution to allow visualization of the dendritic arbor. Pipette solution was kept on ice and shielded from light during the experiment to prevent degradation of the dye or ATP and GTP salts. Patch pipettes had a tip resistance of 4-6MOhm and were mounted in a motorized manipulator (Luigs & Neumann). Liquid junction potential was not corrected. Fast and slow capacitances were compensated, and series resistance was partially compensated (50-80%). Cell health was monitored through the consistency of input current and by calculating series and input resistances with test pulses throughout the recording (fig. S3B). Cells were rejected if any value deviated ±20% of baseline for more than 1min.

CF inputs were stimulated with 0.2ms step pulses using an electrode connected to an isolated current source (SIU91A, Cygnus Technology) and immersed in a glass pipette filled with ACSF. The cell was held in voltage clamp at −10 to −30mV. The stimulus intensity (0-150nA) and location (2-4 sites in the granule cell layer, 50-150μm from the PC soma, spanning the space sub-adjacent to the dendritic arbor) were systematically varied to search for multiple CF inputs. CF excitatory post-synaptic currents (EPSCs), particularly of reduced amplitude in some multi-innervated cells, were distinguished from parallel fiber (PF)-EPSCs by their paired pulse depression (400ms interval) and stable amplitude with small changes in stimulus intensity. PF-EPSCs exhibit paired pulse facilitation and linear amplitude relationship with even small stimulus intensity changes due to the recruitment of additional fibers.

CF-EPSCs were recorded after 20min post-patch to allow sufficient perfusion of cesium and dye. CF multi-innervation was only determined with multiple, discrete and consistent EPSC amplitude steps during both increasing and then decreasing stimulus intensity. After each recording, confocal z-stack images of each cell were obtained with a 63x objective (EC Plan-Neofluar 1.3NA, water immersion, Zeiss) using Zen software.

### Two-Photon Ca^2+^ Imaging

#### Cranial window and GCaMP injection surgeries

Surgeries were performed on animals aged 10-12 weeks under ketamine/xylazine anesthesia (100 and 10mg/kg) with subcutaneous injections of meloxicam (1-2 mg/kg), buprenorphine (0.1 mg/kg), and sterile saline (0.5-1mL) as above. Body temperature was maintained at 35-37L with a feedback dependent heating pad. The skin above the posterior skull was excised and the bone cleaned to implant a metal headframe over the interparietal bone via dental cement. After 3-4 days of recovery, mice were anesthetized and a 4mm craniotomy and durectomy was made at 2.5mm lateral from midline and 2.5mm caudal from lambda, exposing cerebellar simplex, crus I, and anterior crus II. A glass microelectrode with ∼300μm tip diameter was used to inject a viral cocktail with low titer PC-specific L7-Cre (0.5%, AAV1.sL7.Cre.HA.WPRE.hGH.pA; Princeton Neuroscience Institute (PNI) Viral Core Facility; acquired from the lab of Dr. Samuel Wang, Princeton University) and high titer Cre-dependent GCaMP6f (20%, AAV.CAG.Flex.GCaMP6f.WPRE.SV40; Addgene, #100835) was injected ∼300μm below the pial surface of medial and/or lateral crus I (∼900nL per site, 5min wait before needle retraction) and a two-layer cranial window (4mm inner window, Tower Optical; 5mm outer window, Warner Instruments) was implanted over the craniotomy and sealed with dental cement (Metabond).

#### Habituation

The mice recovered for 7 days before habituation began. During the first week, habituation sessions were conducted every other day and consisted of exposure to handling and then to the imaging apparatus and head fixation on the treadmill. In the last 3 days before the experiment (6-10 days after habituation began), mice were habituated every day to head fixation, noises and activity typical during an experimental session, and occasional exposure to multisensory stimuli. Habituation allowed animals to exhibit relative comfort and reduced running behavior.

#### Imaging protocols

Imaging experiments were performed when the GCaMP6f indicator reached stable expression in a sparse cell population (11-20 days post-injection). PC dendrites were imaged at either 61.8 or 30.9Hz using a laser scanning two-photon microscope (Mai Tai DeepSee, Spectra-Physics) with an 8KHz resonant scanning module (Thorlabs) and 16x water immersion objective (Nikon LWD 0.8NA, 3mm WD) controlled with Scanbox (Neurolabware). A digital 4x magnification was used for imaging lateral crus I during spontaneous and multisensory experiments and 2x for imaging medial crus I during whisker stimulation experiments. GCaMP6f was excited using a 920nm femtosecond-pulsed two-photon laser (∼30mW laser power at sample; Spectra-Physics) and fluorescence emission collected by a GAsP PMT (Hamamatsu). Interlocking light shields were fit around the headframe and objective to block ambient light from increasing background noise and to prevent an artifact from blue light stimuli directed at the eye. The microscope is custom designed with a rotating objective turret such that the angle of imaging could be adjusted to capture a perpendicular cross-section of PC dendritic arbors, thus reducing each cell’s imaging profile to reduce the chance of contamination.

In order to increase the imaging rate to 62Hz for spontaneous and multisensory experiments, a narrow field of view was used (656 x 256 pixels scanned instead of 656 x 512 lines for whisker experiments). To image the complete arbor of several PCs when the short axis of the field of view was only ∼153μm, we installed the treadmill, camera, and stimulus apparatuses on a large rotating platform (Thorlabs) such that the animal and all experimental components could be rotated under the objective until the parasagittal plane of PC dendrites aligned with the long axis (∼392μm) of the image. Having the entire width of the dendrites aligned with the scanning direction also provided the benefit of technically optimizing the scanning time for each cell, thus reducing the chance for movement artifacts to appear as a branch-specific signal.

#### Volumetric imaging to confirm morphology

At the end of each imaging session, a volumetric scan was performed over the field of view at the maximum z-resolution of 2μm per step. For each scan, the laser power was turned up to 4-15%, the PMT gain down to 0.7-0.85, and 20-30 images were collected and averaged per step for optimal spatial resolution and morphological detail. Cells were only accepted for use if the somatic and dendritic compartments were entirely visible and major branch points of the primary dendrite were differentiable. These rules restricted our analyses to cells wherein the following parameters were unambiguous: 1) distance from the soma to primary dendrite split, 2) presence or absence of multiple primary trunks emerging from the soma, 3) rostral-caudal distance between branch centroids, and 4) maximal rostral-caudal spread of the whole dendritic arbor. For high-magnification recordings at 4x zoom, where multiple regions of interest (ROIs) were drawn for each major dendritic branch (spontaneous and multisensory experiments), we additionally required the unambiguous distinction of lesser branch points generating sub-compartments.

### Stimulus Conditions

#### Multisensory stimuli

During each experiment, calcium activity was monitored in ∼1-10 cells per animal during 20s imaging sessions. One of eight stimulus types (1. Light, 2. Air Puff, 3. Sound, 4. Light + Puff, 5. Light + Sound, 6. Puff + Sound, 7. Light + Puff + Sound, and 8. Control without stimulus) was triggered 10s after scanning initiation and lasted for 30ms. Light stimulus was a 488nm LED light (Prizmatix) targeted to the ipsilateral eye, Air Puff was delivered at 10psi (Picospritzer III, Parker Hannifin) via a 0.86mm diameter capillary tube positioned 2-3mm from the center of the ipsilateral whisker pad, and Sound stimulus was a 12kHz pure tone produced by speakers (Harman/Kardon) positioned bilaterally at ∼70-80dB. The stimuli were applied with inter-stimulus intervals ≥ 30s. An Arduino Uno microcontroller triggered by the imaging software provided distinct stimulus type triggering output to the light, puff, and tone instruments. The microcontroller was programmed to cycle through stimulus types randomly until 15 trials were acquired of each type (120 trials total).

#### Spontaneous activity

Spontaneous activity was obtained either on a day without sensory stimulation, or from the 10s pre-stimulus baseline period of multisensory imaging sessions.

#### Single whisker stimulation

We obtained a sparse expression of GCaMP6f in PC dendrites and habituated mice to head fixation as described above. In some cases, we used the same mice for this experiment after spontaneous recordings were collected. Before the experiment began, we identified whisker responsive areas of medial Crus I by gently brushing varying numbers of whiskers and observing cellular activity in real time. We then sedated the animals with a minimal dose of ketamine/xylazine (80 and 8mg/kg, respectively) before headfixing them at the two-photon microscope. We strived to conduct the experiment during the 40-60min that the animal was in a stable level of sedation, so whisker responses and spontaneous activity were as close to comparable across whiskers and animals as possible. In a few cases a supplemental dose was required; the experiment was paused while the animal was waking, receiving the supplement, and returning to an equivalent state of sedation.

When the animal reached a state of anesthesia where it stopped actively whisking, a glass capillary tube attached to a rotating motor (SG92R Micro servo, Tower Pro) was manually manipulated to capture a single whisker. Previous work has shown that CF-dependent complex spike responses to whisker stimulation are tuned most commonly to dorsal, caudal, and dorso-caudal directions of whisker displacement (42), so the rotating servo was oriented such that rotation moved the whisker at 135° in the dorsal-caudal direction (fig. S6A).

It is important to note that these experiments are meant as a proof of principle which cannot be construed to represent naturalistic behavior in the awake animal. As previously reported, responses to whisker stimulation, particularly of a single whisker, are very sparse in the anesthetized animal (∼10% of trials). Several approaches could be used to compensate for this. Most obviously, experiments could be conducted in the awake animal when responsiveness is elevated. This approach did not work for us as it is virtually impossible to isolate a single whisker in a capillary tube while the mouse is awake (even if all but some whiskers are trimmed, which we prefer to avoid), and it would be extremely challenging to segregate active whisking from experimental passive whisker deflection. Second, while the animal is under anesthesia numerous individual trials (perhaps 50) with distant stimulus times could be conducted on a single whisker to confidently identify a response that is distinct from spontaneous CF activity. While this approach is more attractive for several reasons, it poses a substantial logistical problem as it would require a long time (>30min) to test a single whisker. This substantially limits how many whiskers could be tested under a consistent state of anesthesia, which massively reduces the chances of identifying whisker responses. Given the limitations of these approaches, we designed an experimentally bootstrapped stimulus wherein each whisker was stimulated many times at a high rate. At the expense of a possible change in responsiveness with repeated stimuli, this allowed many attempts to produce a response, while accelerating our recordings so we could test a large set of whiskers in each animal.

The servo was controlled by an Arduino Uno microcontroller programed to execute a sine function (7° maximal rotation forming an arc circumference of ∼1.5cm at the tip of the capillary tube) at a rate of 2Hz for 50s (100 total stimulations of the whisker with 500ms intervals between starting movement initiations). The microcontroller was triggered by the imaging software with a 10s delay-to-start so spontaneous activity was recorded of ahead of each whisker stimulation. Images were thus 60s in duration (10s spontaneous activity and a 50s stimulation epoch), and two trials were conducted per whisker (200 total stimulations with 20s of spontaneous recording). After two trials, the capillary tube was manually withdrawn and moved to another whisker. Across animals, seven whiskers were stimulated in random order (β, γ, C1, C2, C3, B2, D2; occasionally D1 if D2 was inaccessible).

To sample a wider population of cells and increase the chance of observing whisker responses, imaging was conducted at 2x digital magnification and 31Hz, rather than 4x and 62Hz as above. As in previous 2-photon experiments, a z-stack was obtained at the end of recording to measure morphological properties of each cell and allow ROIs to be drawn manually across dendrites.

### Two-photon image processing

Images were converted to tiffs and motion corrected using custom MATLAB scripts. Cellular ROIs were drawn manually in ImageJ based on volumetric cell reconstructions. Another MATLAB script measured the pixel intensity of each ROI across frames and videos and saved the data as a .mat file. An interactive MATLAB GUI was used to manually confirm detection quality and consistency across imaging sessions to either include or exclude each cell for downstream analysis. Analyses were performed using MATLAB scripts and output for final data shaping, plotting, and statistics in R.

#### Manual event curation

In preliminary experiments, an interactive MATLAB GUI was used to manually curate a findpeaks autodetection of events. Curation involved adjusting rise and peak times as needed and, in cases of a branch-specific event, marking trace locations where a peak was missing. Thus, missed events could be tallied to obtain the number of local events (fig. S5A).

#### Calcium peak detection and comparing inter-branch signals

Raw signal from all ROIs was imported to a custom MATLAB script that performed a five-frame moving window smoothing function and a background correction function. Then, ROI traces were input to the MATLAB version of OASIS deconvolution to obtain times and amplitudes of calcium peaks exceeding 3SD of the baseline. We decided not to distinguish between multiple tightly clustered events producing a single, accumulated large amplitude peak. While accumulated peaks from clustered inputs often retain multiple peaks (a partial peak within the rising phase of the larger event), the slow time constant of the GCaMP6f indicator and the natural variability between small, branch ROIs can alter the appearance of multiple peaks and produce varying spike deconvolutions. This reduces our confidence in the ability to appropriately determine if there is a branch-specific event within a cluster of global events. As such, we identified peak times <4 frames apart – having only 1-2 frames (16-32ms) between detected peaks, which is below the ∼50ms rise time constant of GCaMP6f (*57*) – and took only the second and highest peak or the last in a sequence of >2 events all of which are <4 frames apart.

To compare the deconvolved signal of each branch within a cell, we segregated the data for each trace into five groups: all signal in branch 1, all signal in branch 2, only global events, only local events in branch 1, and only local events in branch 2. Branch number assignment was arbitrary. Subsequent analyses were then performed on each subset of detected peaks individually.

#### Whisker movement traces and timing

First, whisker stimulation times were obtained from 30Hz video recordings (Genie Dalsa, Phase 1 Technology) of the mouse face where the stimulated whisker and the capillary tube moving the whisker were clearly visible. An ROI was drawn in ImageJ at the location in the capillary tube’s movement trajectory where the whisker started to be bent or translated. Thus, when the whisker was moved, the bright capillary tube passed through the ROI and created a time locked peak in light intensity. The entire trace for each video (1min) was extracted in ImageJ for downstream analyses in a custom MATLAB script.

The first derivative of pixel intensity across frames was calculated for each trace, a baseline was measured during the 10s spontaneous period where there was no whisker movement, and whisker stimulus onset times were thus identified as the n-1 frame where a peak in the first derivative exceeded 3SD of the baseline. This also captured the return movement of the capillary tube that returned the whisker to its natural position.

#### Whisker experiment calcium signal

The same methods as described above were used to obtain calcium signal traces from two-photon recordings. Raw signal from all ROIs was imported to a custom MATLAB script that performed the same smoothing and background correction as described above. Then, ROI signals for each cell were analyzed two ways: averaged into a whole cell signal trace or kept separate to independently assess each branch. We input either whole cell or branch traces into OASIS deconvolution to obtain times and amplitudes of calcium peaks exceeding 2SD of the baseline and >10% the amplitude of the largest detected peak. These parameters allowed the initial detection of smaller and less typically shaped events and the post-hoc elimination of excessively small events that could be noise during a period of elevated baseline or a highly irregular peak possibly due to a PF burst.

As above, when comparing branches within a cell, we segregated deconvolved calcium events into the same five groups. The next step, determining if the events constituted a response to the stimulated whisker, was then performed on each subset of detected peaks individually.

As two 1min trials were conducted for each whisker, we concatenated the event amplitude and timing data from each trial of the same whisker. We then compared the event rise times (when the rising phase began) with the whisker stimulus times to assess how many peaks occurred during 150ms (5 frame) time windows after each whisker movement, as opposed to non-response windows of 150ms before each whisker movement or the 10s spontaneous time window when there was no whisker stimulation. Probabilities and amplitudes of response and non-response events were thus calculated for each ROI-averaged whole cell and individual branch ROIs. Since whisker responses are known to be very sparse under anesthesia, we stimulated each whisker 100 times per trial (for a total of 200 trials over 2min) to experimentally bootstrap response probabilities. The repetition allowed us to calculate not only the absolute probability of response, but the variability of the response and non-response probabilities across frames relative to whisker movement time such that we could obtain a Z-score of the response probability using the following formula:

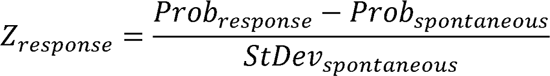

For each set of input event times, if the Z_response_ > 1.96 (2SD) – high enough to reject H_0_ with α < 0.05 – then we considered this a response to the whisker.

### Statistics and Quantifications

Statistical analysis was carried out using R (v4.2.1). Data following a normal distribution was tested with unpaired two-tailed Student’s t-tests to compare two groups or ANOVA followed by a Tukey post-hoc correction for repeated measures to compare more than two groups and/or multiple factors. A one sample Student’s t-test was used to compare individual groups with a specific, benchmark value where appropriate. Otherwise, for non-parameteric data, a Wilcoxon rank sum test was used for two group comparisons. For comparisons of contingency tables from data with nominal categories, a Pearson’s Chi-squared test was used. To assess a relationship between two continuous variables, we used a linear regression and report the adjusted R^2^ value and significance. For all analyses, α = 0.05 was used to determine significance and figure panels refer to the significance of comparisons in the following way: ns p > 0.05, * p ≤ 0.05, ** p ≤ 0.01, *** p ≤ 0.001.

### Main Figures

**Fig 1C**: Pearson’s Chi-squared, Ratio of cell morphologies ∼ Lobule; n = 6,646 cells, p < 0.001, X2 = 170.18.

**Fig 1F**: Pearson’s Chi-squared, Ratio of cell morphologies ∼ Lobule; n = 1,350 cells, p < 0.001, X2 = 106.83.

**Fig 1G**: Morphology and Species: Two-way ANOVA, Morphology p < 0.001, Species p = 0.937; Tukey’s HSD post-hoc: Normative, n = 30 and 20 lobules, 46.53 ± 2.2 and 5.82 ± 1.42 Percent of cells (%), Normative_Mouse_ vs. Normative_Human_, p < 0.001; Split, n = 30 and 21 lobules, 36.6 ± 2.04 and 47.7 ± 2.4 Percent of cells (%), Split_Mouse_ vs. Split_Human_, p = 0.013; Poly, n = 29 and 21 lobules, 17.46 ± 2.45 and 46.76 ± 2.97 Percent of cells (%), Poly_Mouse_ vs. Poly_Human_, p < 0.001; Normative_Mouse_ vs. Split_Mouse_, p = 0.015; Normative_Mouse_ vs. Poly_Mouse_, p < 0.001; Split_Mouse_ vs. Poly_Mouse_, p < 0.001; Normative_Human_ vs. Split_Human_, p < 0.001; Normative_Human_ vs. Poly_Human_, p < 0.001; Split_Human_ vs. Poly_Human_, p = 1.

**Fig 2D**: Pearson’s Chi-squared, Number of climbing fibers ∼ Morphology; n = 159 cells, p = 0.003, χ^2^ = 15.71.

**Fig 2E**: 1 vs. 2+ CFs: n = 135 and 24 cells, −763.65 ± 2.51 and −978.75 ± 13.45 CF EPSC amplitude (pA), p = 0.006, Two-tailed Student’s t-test; Weak CFs: n = 24 cells, −389.13 ± 10.3 CF EPSC amplitude (pA), p < 0.001, One-way Student’s t-test, µ = 0.

**Fig 2F**: 1 vs. 2+ CFs, all PCs: n = 135 and 24 cells, 43.74 ± 3.13 and 16.86 ± 3.28 Split distance (µm), p < 0.001, Two-tailed Student’s t-test.

**Fig 2G**: 1 vs. 2+ CFs, mono-planar PCs: n = 85 and 9 cells, 54.48 ± 3.26 and 87.09 ± 7.73 Branch distance (µm), p = 0.002, Two-tailed Student’s t-test.

**Fig 2H**: 1 vs. 2+ CFs, Poly PCs: n = 26 and 8 cells, 67.32 ± 5.7 and 95.91 ± 10 Trunk angle (°), p = 0.029, Two-tailed Student’s t-test.

**Fig 3F**: Global vs. Local events: n = 95 cells, 0.29 ± 0.009 and 0.19 ± 0.007 Mean amplitude (ΔF/F0), p < 0.001, Two-tailed Student’s t-test.

**Fig 3G**: Normative, Split, Poly: One-way ANOVA, p < 0.001; Tukey’s HSD post-hoc, n = 32, 55, 8 cells, 17.36 ± 2.12, 36.62 ± 2.54, and 51.01 ± 9.76 Local events (%); Normative vs. Split, p < 0.001; Normative vs. Poly, p < 0.001; Split vs. Poly, p = 0.087.

**Fig 3H**: Normative, Split, Poly: Kruskal-Wallis test, p < 0.001; Pairwise Wilcoxon rank sum test, n = 32, 55, 8 cells, 0.37 ± 0.03, 0.24 ± 0.03, and 0.079 ± 0.04 Adjusted R^2^; Normative vs. Split, p = 0.002; Normative vs. Poly, p < 0.001; Split vs. Poly, p = 0.016.

**Fig 3I**: Pearson’s Chi-squared, Rate of significant branch covariation ∼ Morphology; n = 95 cells, p = 0.007, χ^2^ = 9.8.

**Fig 3J**: Linear Regression, Inter-branch amplitude scale (sd) ∼ Split distance (µm); n = 105 cells, p < 0.001, R^2^ = 0.099.

**Fig 3K**: Linear Regression, excluding multi-planar cells, Inter-branch amplitude scale (sd) ∼ Dendrite width (µm); n = 109 cells, p = 0.002, R^2^ = 0.078.

**Fig 4C**: Normative, Split, Poly: One-way ANOVA, p < 0.001; Tukey’s HSD post-hoc, n = 33, 112, 24 cells, 2.72 ± 0.36, 4.48 ± 0.29, and 5.78 ± 0.39 Mean local responses (#); Normative vs. Split, p = 0.004; Normative vs. Poly, p < 0.001; Split vs. Poly, p = 0.089.

**Fig 4D**: Normative, Split, Poly: One-way ANOVA, p = 0.02; Tukey’s HSD post-hoc, n = 33, 112, 24 cells, 4.76 ± 0.42, 6.29 ± 0.33, and 6.96 ± 0.56 Inter-whisker difference (#); Normative vs. Split, p = 0.041; Normative vs. Poly, p = 0.029; Split vs. Poly, p = 0.623.

**Fig 4F**: Pearson’s Chi-squared, Response category ∼ Morphology; n = 169 cells, p = 0.035, χ^2^ = 6.73.

**Fig 4G**: Unresponsive, Global, and Lateral: One-way ANOVA, p = 0.02; Tukey’s HSD post-hoc, n = 75, 42, 52 cells, 27.35 ± 2.06, 28.24 ± 3.58, and 18.73 ± 2.05 Split distance (µm); Unresponsive vs. Global, p = 0.967; Unresponsive vs. Lateral, p = 0.034; Global vs. Lateral, p = 0.043.

**Fig 4H**: Linear Regression, Local events (%) ∼ Receptive field (whiskers); n = 151 cells, p = 0.005, R^2^ = 0.046.

**Fig 5C**: Morphology and Stimulus Category: Two-way ANOVA, Morphology p < 0.001, Stimulus Category p = 0.047; Tukey’s HSD post-hoc: Uni-modal, n = 24 and 38 cells, 18.06 ± 1.94 and 31.23 ± 2.98 Maximum local events (%), Uni_Normative_ vs. Uni_Split+Poly_, p = 0.014; Multi-modal, n = 24 and 38 cells, 21.11 ± 3.02 and 38.95 ± 3.01 Maximum local events (%), Multi_Normative_ vs. Multi_Split+Poly_, p < 0.001; Uni_Normative_ vs. Multi_Normative_, p = 0.917; Uni_Split+Poly_ vs. Multi_Split+Poly_, p = 0.176.

**Fig 5F**: Normative vs. Split+Poly PCs: n = 24 and 38 cells, 29.84 ± 3.51 and 45.58 ± 4.43 ΔBranch response range (%), p = 0.008, Two-tailed Student’s t-test.

**Fig 5G**: Normative vs. Split+Poly PCs: n = 24 and 38 cells, −1.64 ± 1.44 and −1.57 ± 3.57 ΔBranch response mean (%), p = 0.986, Two-tailed Student’s t-test.

**Fig 5H**: Normative vs. Split+Poly PCs: n = 24 and 38 cells, 31.47 ± 4.02 and 47.14 ± 5.55 ΔBranch response bilaterality (%), p = 0.027, Two-tailed Student’s t-test.

### Supplementary Figures

**fig S3C**: Pearson’s Chi-squared, Multi-CF rate (%) ∼ Split distance (µm); n = 159 cells, p = 0.025, χ^2^ = 18.65.

**fig S3D**: 1 vs. 2+ CFs, non-Poly PCs excluding Normative PCs with no split: n = 79 and 16 cells, 36.77 ± 2.17 and 25.29 ± 3.25 Split distance (µm), p = 0.006, Two-tailed Student’s t-test **fig S3E**: 1 vs. 2+ CFs, non-Poly PCs: n = 79 and 16 cells, 96.97 ± 2.61 and 105.44 ± 4.95 Split angle (°), p = 0.16, Two-tailed Student’s t-test.

**fig S3F**: 1 vs. 2+ CFs, Poly PCs: n = 26 and 8 cells, 67.32 ± 5.7 and 95.91 ± 10 Trunk angle (°), p = 0.029, Two-tailed Student’s t-test.

**fig S3G**: 1 vs. 2+ CFs, all PCs excluding Normative PCs with no split: n = 104 and 24 cells, 66.85 ± 2.41 and 83.19 ± 6.05 Arbor separation (µm), p = 0.018, Two-tailed Student’s t-test

**fig S3H**: 1 vs. 2+ CFs, Poly PCs: n = 26 and 8 cells, 21.23 ± 0.35 and 21.2 ± 0.49 Soma diameter (µm), p = 0.97, Two-tailed Student’s t-test.

**fig S3M**: Linear Regression, CF EPSC amplitude (nA) ∼ Age (days); n = 24 cells, p < 0.001, R^2^ = 0.445.

**fig S3N**: Dominant CF to multi-CF PCs: Linear Regression, CF EPSC amplitude (nA) ∼ Age (days); n = 24 cells, p = 0.235, R^2^ = 0.021; Single CF to mono-CF PCs: Linear Regression, CF EPSC amplitude (nA) ∼ Age (days); n = 126 cells, p = 0.383, R^2^ = −0.002.

**fig S3O**: Normative, Split, and Poly: Kruskal-Wallis rank sum test, p = 0.21; Split vs. Poly PCs: n = 15 and 8 cells, 36.48 ± 1.23 and 53.96 ± 3.16 Small CF : big CF EPSC amplitude (%), p = 0.112, Two-tailed Student’s t-test (p = 0.056, One-tailed Student’s t-test).

**fig S3P**: Gyrus, Bank, and Sulcus: One-way ANOVA, p = 0.058; Tukey’s HSD post-hoc, n = 6, 11, 7 cells, 27.91 ± 9.32, 40.69 ± 5.78, and 56.16 ± 7.04 Small CF : big CF EPSC amplitude (%); Gyrus vs. Bank, p = 0.431; Gyrus vs. Sulcus, p = 0.048; Bank vs. Sulcus, p = 0.27.

**fig S3Q**: Linear Regression, Small CF : big CF EPSC amplitude (%) ∼ Trunk angle (°); n = 8 cells, p = 0.007, R^2^ = 0.687.

**fig S3R**: Linear Regression, Small CF : big CF EPSC amplitude (%) ∼ Split distance (µm); n = 23 cells, p = 0.138, R^2^ = 0.059.

**fig S4L**: 1 vs. 2+ CFs, all PCs: n = 135 and 24 cells, 37.48 ± 0.79 and 42.58 ± 2.54 Age (days), p = 0.065, Two-tailed Student’s t-test.

**fig S5A**: Normative, Split, Poly: One-way ANOVA, p = 0.028; Tukey’s HSD post-hoc, n = 25, 29, 5 cells, 5.27 ± 1.18, 12.7 ± 2.58, and 16.74 ± 7.25 Local events (%); Normative vs. Split, p = 0.05; Normative vs. Poly, p = 0.11; Split vs. Poly, p = 0.75.

**fig S5B**: Normative, Split, Poly: One-way ANOVA, p = 0.01; Tukey’s HSD post-hoc, n = 25, 29, 5 cells, 0.006 ± 0.07, 0.351 ± 0.08, and 0.19 ± 0.17 Ca^2+^ amplitude coefficient of variation between vs. within branches (CV); Normative vs. Split, p = 0.006; Normative vs. Poly, p = 0.71; Split vs. Poly, p = 0.77.

**fig S5C**: Linear Regression, Amplitude scale (sd) ∼ Compartment separation (µm); n = 95 cells, p = 0.017, R^2^ = 0.05.

**fig S5D**: Normative, Split, Poly: One-way ANOVA, p = 0.389; Tukey’s HSD post-hoc, n = 32, 55, 8 cells, 1.26 ± 0.04, 1.38 ± 0.06, and 1.37 ± 0.17 Event rate (Hz); Normative vs. Split, p = 0.366; Normative vs. Poly, p = 0.757; Split vs. Poly, p = 0.998.

**fig S5E**: Split vs. Normative PCs: n = 13 animals, 102.72 ± 5.76 Split / Normative Rate (%), p = 0.645, One-way Student’s t-test, µ = 100.

**fig S5F**: Linear Regression, Split / Normative Rate (%) ∼ Local gap (%); n = 13 animals, p = 0.813, R^2^ = −0.085.

**fig S5G**: Linear Regression, Rate above minimum (Hz) ∼ Local events (%); n = 94 cells, p = 0.001, R^2^ = 0.1.

**fig S5H**: Linear Regression, Event rate (Hz) ∼ Local events (%); n = 95 cells, p = 0.008, R^2^ = 0.06.

**fig S5I**: Normative, Split, Poly: One-way ANOVA, p = 0.043; Tukey’s HSD post-hoc, n = 32, 55, 8 cells, 2.23± 0.09, 2.74 ± 0.14, and 2.71 ± 0.46 Event rate (Hz); Normative vs. Split, p = 0.036; Normative vs. Poly, p = 0.387; Split vs. Poly, p = 0.995.

**fig S5J**: Split vs. Normative PCs: n = 13 animals, 126.25 ± 8.29 Split / Normative Rate (%), p = 0.008, One-way Student’s t-test, µ = 100.

**fig S5K**: Linear Regression, Split / Normative Rate (%) ∼ Local gap (%); n = 13 animals, p = 0.005, R^2^ = 0.48.

**fig S5L**: Linear Regression, Rate above minimum (Hz) ∼ Local events (%); n = 94 cells, p < 0.001, R^2^ = 0.14.

**fig S5M**: Linear Regression, Event rate (Hz) ∼ Local events (%); n = 95 cells, p < 0.001, R^2^ = 0.26.

**fig S5N**: Normative, Split, Poly: One-way ANOVA, p = 0.019; Tukey’s HSD post-hoc, n = 13, 16, 6 animals, 2.12 ± 0.23, 3.36 ± 0.47, and 2.85 ± 0.31 Event rate gap (Hz); Normative vs. Split, p = 0.014; Normative vs. Poly, p = 0.384; Split vs. Poly, p = 0.607.

**fig S5O**: Linear Regression, Event rate gap (Hz) ∼ Local events (%); n = 95 cells, p < 0.001, R^2^ = 0.2.

**fig S5P**: Minimum vs. non-minimum cells: n = 16 and 78 cells, 13.39 ± 4.93 and 22.21 ± 2.11 Local events (%), p = 0.032, Wilcoxon rank sum test.

**fig S6E**: Normative, Split, Poly: One-way ANOVA, p = 0.002; Tukey’s HSD post-hoc, n = 28, 99, 22 cells, 22.32 ± 3.58, 36.88 ± 2.34, and 43.36 ± 4.29 Local events (%); Normative vs. Split, p = 0.007; Normative vs. Poly, p = 0.003; Split vs. Poly, p = 0.428.

**fig S6F**: Unresponsive, Global, and Lateral: One-way ANOVA, p < 0.001; Tukey’s HSD post-hoc, n = 75, 42, 52 cells, 32.13 ± 2.62, 32.72 ± 3.24, and 46.76 ± 2.51 Local events (%); Unresponsive vs. Global, p = 0.988; Unresponsive vs. Lateral, p < 0.001; Global vs. Lateral, p = 0.004.

**fig S7B**: Pearson’s Chi-squared, Response Type ∼ Morphology; n = 2,520 and 3,990 events, 24 and 38 cells, p < 0.001, χ^2^ = 169.13.

**fig S7C**: Morphology and Stimulus Category: Two-way ANOVA, Morphology p = 0.88, Stimulus Category p = 0.002; Tukey’s HSD post-hoc: Uni-modal, n = 24 and 38 cells, 70.37 ± 4.29 and 70.12 ± 4.03 Response probability (%), Uni_Normative_ vs. Uni_Split+Poly_, p = 0.999; Multi-modal, n = 24 and 38 cells, 82.22 ± 4.83 and 83.73 ± 3.53 Response probability (%), Multi_Normative_ vs. Multi_Split+Poly_, p = 0.994; Uni_Normative_ vs. Multi_Normative_, p = 0.285; Uni_Split+Poly_ vs. Multi_Split+Poly_, p = 0.053.

**fig S7D**: Morphology and Stimulus Category: Two-way ANOVA, Morphology p < 0.001, Stimulus Category p < 0.001; Tukey’s HSD post-hoc: Control, n = 24 and 38 cells, 5 ± 1.15 and 7.54 ± 1.38 Maximum local events (%), Ctrl_Normative_ vs. Ctrl_Split+Poly_, p = 0.983; Uni-modal, n = 24 and 38 cells, 18.06 ± 1.94 and 31.23 ± 2.98 Maximum local events (%), Uni_Normative_ vs. Uni_Split+Poly_, p = 0.014; Multi-modal, n = 24 and 38 cells, 21.11 ± 3.02 and 38.95 ± 3.01 Maximum local events (%), Multi_Normative_ vs. Multi_Split+Poly_, p < 0.001; Ctrl_Normative_ vs. Uni_Normative_, p = 0.019; Ctrl_Split+Poly_ vs. Uni_Split+Poly_, p < 0.001 ; Uni_Normative_ vs. Multi_Normative_, p = 0.975; Uni_Split+Poly_ vs. Multi_Split+Poly_, p = 0.166.

**fig S7E**: Morphology and Stimulus Category: Two-way ANOVA, Morphology p < 0.001, Stimulus Category p < 0.001; Tukey’s HSD post-hoc: Control, n = 24 and 38 cells, 3.33 ± 0.8 and 4.74 ± 0.9 Directional maximum local events (%), Ctrl_Normative_ vs. Ctrl_Split+Poly_, p = 0.997; Uni-modal, n = 24 and 38 cells, 13.61 ± 1.73 and 23.33 ± 2.84 Directional maximum local events (%), Uni_Normative_ vs. Uni_Split+Poly_, p = 0.029; Multi-modal, n = 24 and 38 cells, 13.61 ± 1.77 and 26.14 ± 2.7 Directional maximum local events (%), Multi_Normative_ vs. Multi_Split+Poly_, p = 0.001; Ctrl_Normative_ vs. Uni_Normative_, p = 0.043; Ctrl_Split+Poly_ vs. Uni_Split+Poly_, p < 0.001 ; Uni_Normative_ vs. Multi_Normative_, p = 1.0; Uni_Split+Poly_ vs. Multi_Split+Poly_, p = 0.914.

**fig S7F**: Normative PCs: n = 24 cells, 3.06 ± 0.44 Uni vs. Multi-modal difference in maximum local events, p = 0.171, One-way Student’s t-test, µ = 0; Split+Poly PCs: n = 38 cells, 7.72 ± 0.34 Uni vs. Multi-modal difference in maximum local events, p < 0.001, One-way Student’s t-test, µ = 0.

**fig S7I**: Unimodal Normative vs. Split+Poly PCs: n = 24 and 38 cells, 21.41 ± 3.66 and 32.71 ± 4.41 ΔBranch response range (%), p = 0.053, Two-tailed Student’s t-test; Multimodal Normative vs. Split+Poly PCs: n = 24 and 38 cells, 17.29 ± 2.14 and 27.44 ± 3.12 ΔBranch response range (%), p = 0.01, Two-tailed Student’s t-test.

**fig S7J**: Unimodal Normative vs. Split+Poly PCs: n = 24 and 38 cells, −3.18 ± 2.11 and 3.73 ± 4.51 ΔBranch response mean (%), p = 0.172, Two-tailed Student’s t-test; Multimodal Normative vs. Split+Poly PCs: n = 24 and 38 cells, −0.48 ± 1.43 and −4.11 ± 3.35 ΔBranch response mean (%), p = 0.329, Two-tailed Student’s t-test.

**fig S7K**: Unimodal Normative vs. Split+Poly PCs: n = 24 and 38 cells, 24.59 ± 0.44 and 28.98 ± 6.63 ΔBranch response bilaterality (%), p = 0.602, Two-tailed Student’s t-test; Multimodal Normative vs. Split+Poly PCs: n = 24 and 38 cells, 17.78 ± 2.52 and 31.56 ± 4.65 ΔBranch response bilaterality (%), p = 0.013, Two-tailed Student’s t-test.

## Acknowledgments

For valuable advice and technical support, we thank Hansel lab members T.F. Lin, A. Silbaugh, and T. Pham. We thank R.A. Eatock and P. Mason (UChicago Neurobiology) for insightful discussions. For crucial feedback on the manuscript, we thank W. Wei and M. Sheffield (UChicago Neurobiology) as well as S. S. Wang (Princeton). M. Sheffield and S. S. Wang also provided preliminary viral tools. C. Ross (Organismal Biology and Anatomy, OBA; Anatomical Gift Association of Illinois, AGAI), G. Voegele (OBA), W. O’Connor, S. Shimkus, and M. Torres (AGAI) provided human tissue.

## Author contributions

SEB and CH designed the experiments, SEB performed the investigation, formal analysis, visualization, and wrote the original draft, SEB and CH acquired funding and edited the text, CH supervised the work.

## Data and materials availability

All data are available in the manuscript or the supplementary materials

---

**Fig. S1.**
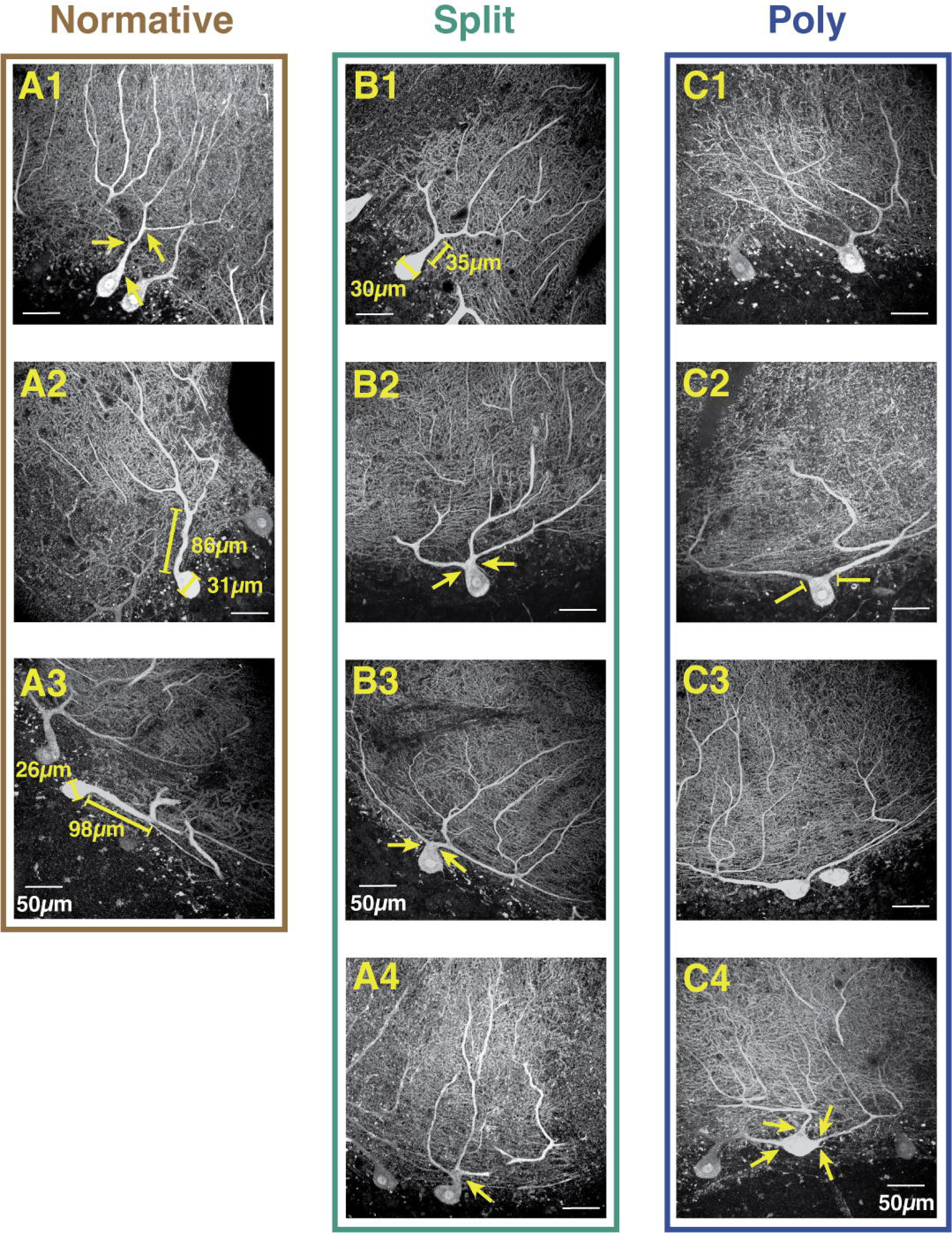
Categorization of human Purkinje cells by proximal primary dendrite geometry. Human PCs were visualized in unembalmed tissue of one individual following immunolabeling for calbindin. (**A1-3**) Normative PCs have one trunk emerging from the soma and either 1) no bifurcation and no proximal minor branches ramifying >200μm laterally (**A1**) or 2) a bifurcation that is >2x the somatic diameter (**A2-3**). We used a threshold bifurcation distance of two somatic diameters measured from the base of the trunk where the circular dimension of the soma ends to the middle of the branch point. As seen in (**A2-3**), the visible bifurcations occur more than 2x the somatic diameter. To accelerate categorization, the distances were only measured when it was not obvious, as it was in most cases, whether the threshold was exceeded. (**B1-3**) Split PCs have one trunk emerging from the soma and either 1) no bifurcation but a large, proximal minor branch ramifying >200μm laterally or 2) a bifurcation that is <2x the somatic diameter (**B1**). Split PCs had spectrums of bifurcation distances and angles. (**B2-3**) The most challenging cases for categorization, which were common, presented with multiple primary dendrites emerging from a compartment which was difficult to define as somatic or dendritic. For our analyses we used the curvature of the space between the branch points and the center of the soma as an indication. As shown in (**B2-3**), there is a minor inward curvature of the superficial somatic compartment, like if the top of the soma was pinched into a single dendritic trunk from which the primary dendrite branches emerge. Compare this with (**C1-2**) where there is no pinching of the superficial soma such that the outer diameter merges directly with the primary dendrites and thus fits a definition of Poly PC. (**B4**) The most challenging cases for categorization, which were rare, presented with proximal bifurcations of the dendrite that were very proximal to the soma, but either asymmetrical or producing more than two primary dendrites. (**C1-4**) Poly PCs have multiple dendritic trunks emerging directly from the somatic compartment. Poly PCs had a spectrum of angles between emerging trunks, from acute (**C1**) or intermediate (**C2**) to emerging from opposite poles of a horizontally oriented somatic compartment. In the latter case, primary dendrites may continue ramifying in opposing directions to form entirely distinct and distant compartments as in (**C3**) or rapidly curve upward to preserve a somewhat compact set of compartments. (**C4**) While less common than two primary dendrites, we also observed many cases where Poly PCs had three or more dendrites emerging at varying relative angles.

**Fig. S2.**
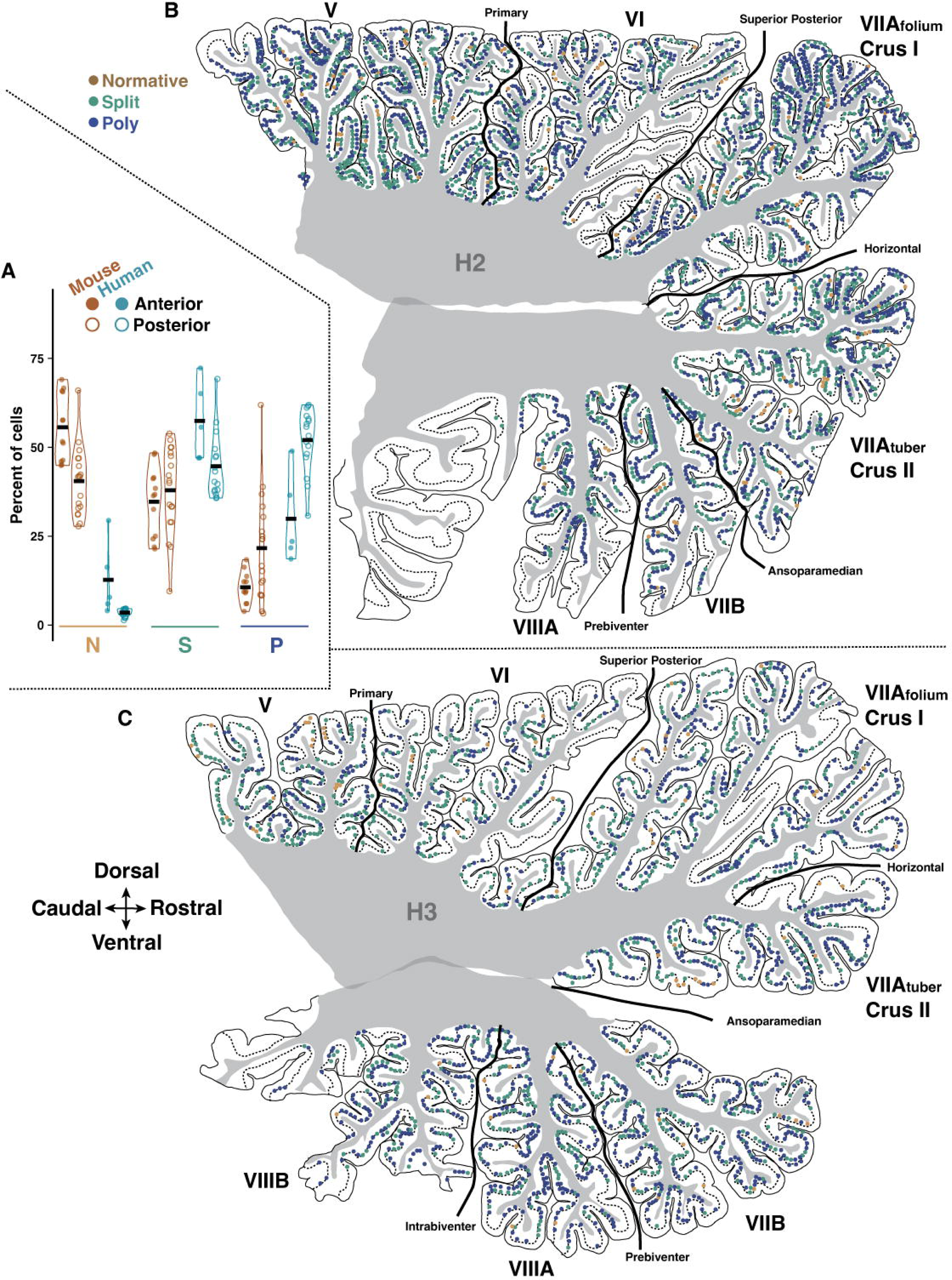
Reconstructed parasagittal sections of human cerebellar hemisphere. (**A**) Summary of relative cell morphology rates across anterior and posterior lobules of human and mouse (see table 3 for statistics). (**B-C**) Cross-section maps of human mid-hemisphere demonstrating the distribution of PC morphological types by region for individual specimens H2 and H3. H4 is presented in (Fig. 1B). See table 1 for quantification.

**Fig. S3.**
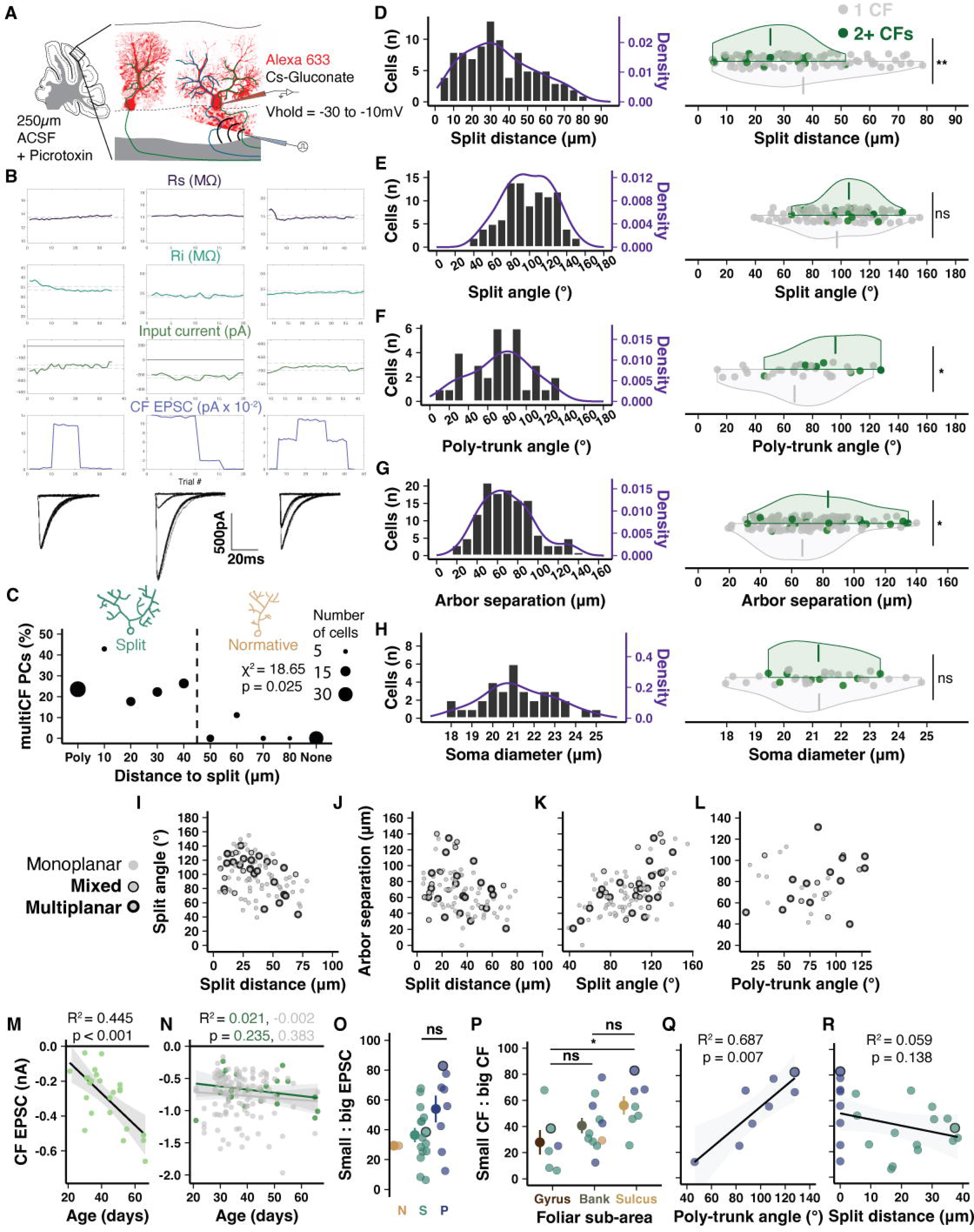
Purkinje dendrite morphology correlates with climbing fiber multi-innervation. (**A**) Schematic of PC whole-cell patch-clamp recordings and CF stimulation in acute cerebellar slices. A cesium internal solution with Alexa 633 dye is used to reduce space clamp error, increase detection of more distant CF inputs, and label the cell for confocal imaging at the end of the experiment. Note, cesium and depolarized holding potentials distort the absolute size of the EPSC. (**B**) Representative physiology and CF-EPSC traces evoked by increasing and decreasing stimulation intensities. *Left*, a mono-innervated PC exhibits only one CF-EPSC amplitude regardless of stimulation intensity above threshold. *Middle*, a multi-innervated PC with discrete CF-EPSC amplitude steps constituting the stimulation of one small CF (∼200pA) and then the summed EPSC amplitude of stimulating two CFs (∼1.2nA) at a higher stimulation intensity. *Right*, discrete CF-EPSC steps indicating that this PC receives two CFs of approximately equal size (∼400-450pA). (**C**) Inverse linear relationship between the rate of multi-innervation and distance from soma to primary dendrite split (n = 50 animals, 159 cells). (**D-H**) Whole population distributions (left) and distributions by CF innervation type (right) of morphological parameters: split distance (n = 79, 16), split angle (n = 79, 16), poly-trunk angle (n = 26, 8), arbor separation (n = 109, 16), and poly PC soma size (n = 26, 8). (**I-L**) Relationships between morphological parameters and Purkinje dendrite planarity. (**M**) Age dependency of weaker CF EPSC amplitude to multi-CF PCs indicates a possible continued development following circuit maturation (n = 24). (N) In contrast, neither the dominant CF on multi-CF PCs (green, n = 24) nor lone CFs of mono-innervated PCs (grey, n = 135) show a relationship between age and EPSC amplitude. (**O**) The ratio of EPSC amplitudes between strong and weak CF inputs to multi-innervated PCs are widely varied across cell morphologies. Bordered points indicate cells with 3 CFs where the smallest of the three inputs is compared with the largest (n = 24 cells). (**P**) Among all multi-innervated PCs, the ratio of EPSC amplitude between the smaller and larger detected CF is smaller in the gyrus (28%) than the sulcus (56%; n = 6, 11, 7 cells). (**Q**) Among multi-innervated Poly PCs, a wider dendrite separation angle correlates with greater EPSC amplitude parity between multiple CFs (n = 8 cells). (**R**) Among multi-innervated PCs, an earlier dendrite split (0 for Poly PCs) does not correlate with greater EPSC amplitude parity (n = 23).

**Fig. S4.**
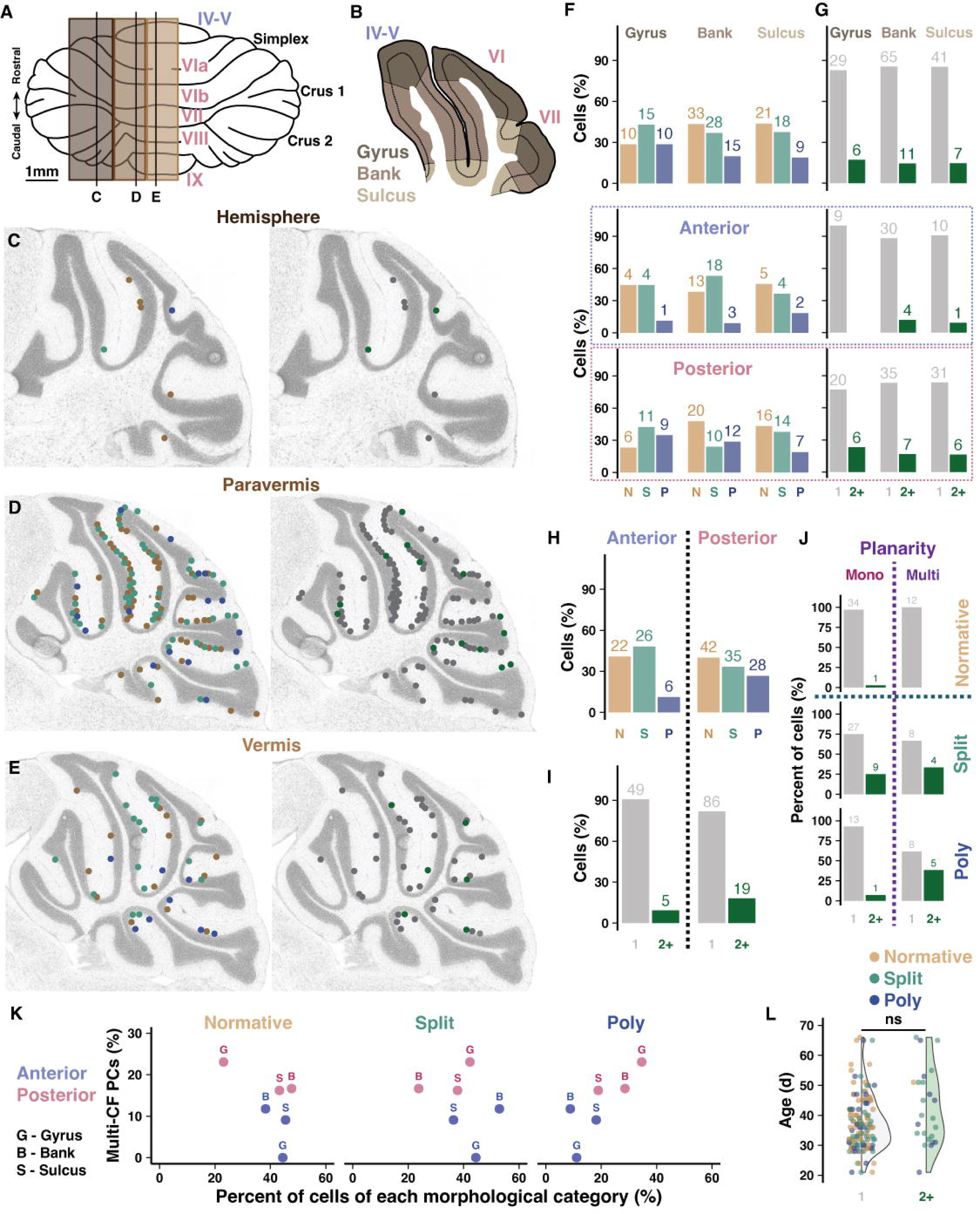
Prevalence of climbing fiber multi-innervation and dendritic morphology across cerebellar regions. (**A**) Schematic of the mouse cerebellum and the medio-lateral location of acute slices used in whole cell patch clamp recordings. (**B**) Example of a sagittal section demonstrating the division of foliar sub-areas. (**C-E**) Slice maps indicating the location of each recorded PC and its morphology (left) or CF innervation pattern (right, multi-CF PCs are green). (**F-G**) Distribution of morphological type (left) or CF innervation pattern (right) across foliar sub-areas for all cells (top) or by lobule region (bottom, anterior vs posterior). Numbers above bars are absolute number of cells. (**H-I**) Distribution of morphological type (**H**) or CF innervation pattern (**I**) by lobule region without sub-dividing foliar sub-area. (**J**) Ratios of multi-innervated PCs by morphology or dendritic planarity. Previously, dendrite planarity has been linked to CF innervation patterns (30), but we find that the presence of multiple primary dendrites – largely requisite to multiplanarity – is slightly more predictive of CF multi-innervation (4 : 96% vs. 55 : 45%). Some PCs could not be clearly identified by planarity, so only a subset of data is presented. (**K**) Relationship between percent of cells with multi-CF innervation across lobule region (anterior in blue, posterior in pink) and foliar sub-area (G – gyrus, B – bank, S – sulcus) and the percent of cells of either Normative, Split, or Poly structure. (**L**) No effect of age on the presence of CF multi-innervation (n = 135 and 24 cells).

**Fig. S5.**
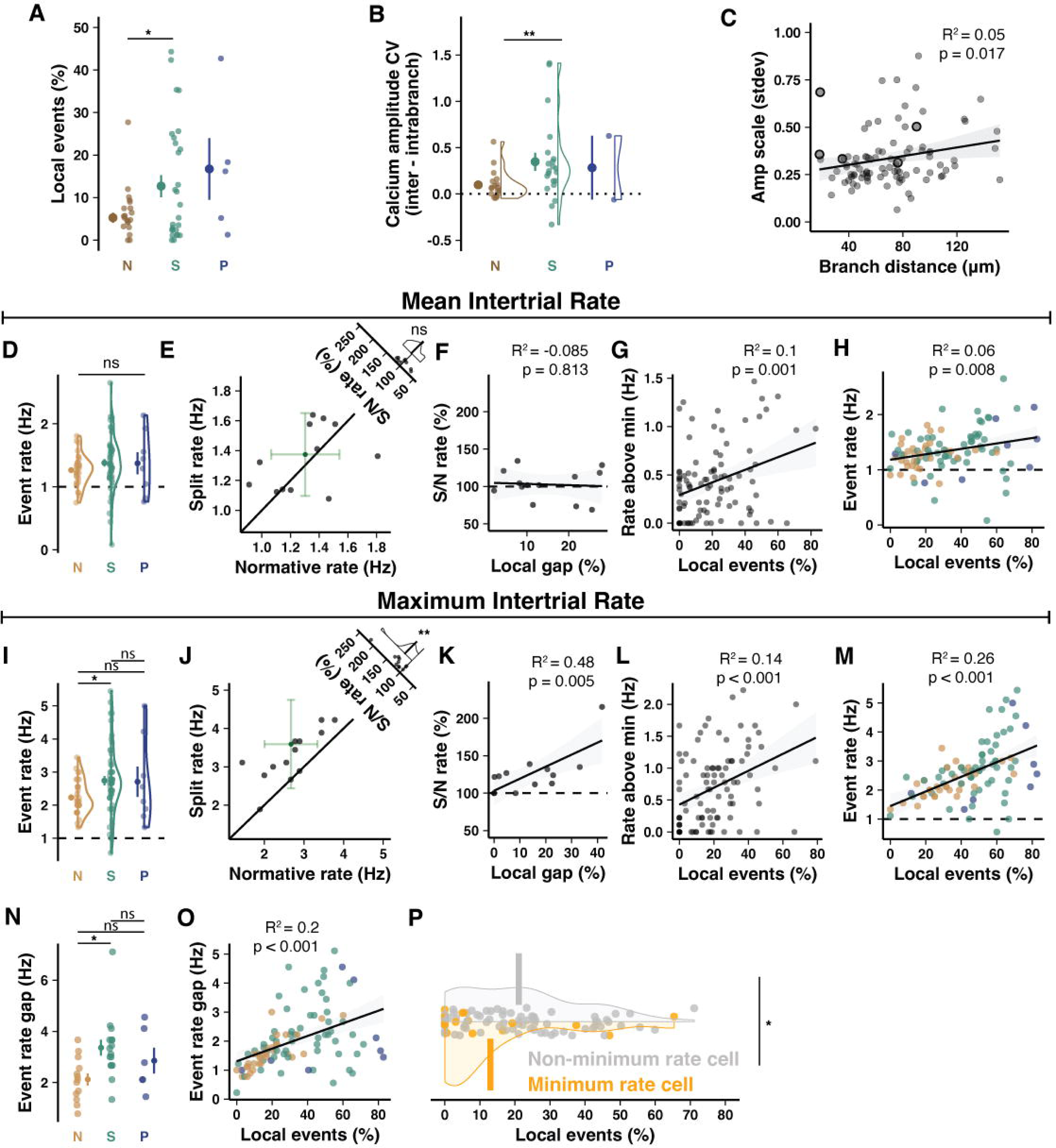
Characterizing calcium signal heterogeneity and subsequent elevation of event rate and total dynamic range. (**A**) Local events identified manually in a subset of data finds a lower baseline frequency of local events but an identical pattern of elevation in Split and Poly PCs (n = 16 animals, n = 25, 29, and 5 cells). (**B**) Analyzing the coefficient of variation of event amplitudes within vs. between branches of each cell shows that, compared to Normative cells, Split PCs have more variability between branches that is not found within branches. This indicates that heterogeneity is a feature of dendritic macro-compartments instead of small sub-branches within a primary dendrite (n = 16 animals, n = 25, 29, and 5 cells). (**C**) Variability of the inter-event amplitude scale (the mean ratio of event amplitude between branches) as a function of the distance between the centroid of dendritic branches (n = 23 animals, n = 95 cells). (**D-H**) Converging CFs likely originate from neighboring IO cells with local synchrony via gap junction coupling and convergence of IO input pathways. Thus, multiple converging CFs likely exhibit some overlapping activity and supply only modest elevations of input frequency. To test this, we compared total CF-dependent event rates to the quantity of heterogeneity. Lateral crus I PCs are largely zebrin(+), but may still have modest variation in zebrin identity and therefore CF input rate (*3*). To compensate, we analyzed raw and normalized inter-trial event rates pooled across PCs and animals. When mean spontaneous event rate is measured during control trials, we made several observations. (**D**) There is no difference between mean event rates by morphology (n = 23 animals, n = 32, 55, and 8 cells). (**E**) To control for differences in CF input rates based on PC molecular patterning identities, we can analyze only cells in animals where Split and Normative cells were both present in a local field of view. There is no bias in the Split relative to Normative PC mean rate within each animal (n = 13 animals). (**F**) Relatedly, there is no relationship between the local event ratio difference between Normative and Split PCs (Local ratio gap) and the ratio of Split to Normative PC event rates (n = 13 animals). (**G**) Controlling for event rate differences across animals by normalizing event rates to the local minimum shows an elevation of event rates, regardless of morphology, in PCs with higher local event ratios (n = 23 animals, n = 94 cells). (**H**) As in J, but without normalizing event rate, PCs with higher local event ratios have modestly elevated event rates (n = 23 animals, n = 95 cells). (**I-M**) Numerous variables could be minimizing the effect size observed by using the mean event rate. On the other hand, analyzing the maximum event rate might provide a more direct indication of the elevation of event rate afforded by putative multi-CF innervation. Thus, we performed the same measurements as in (**D-H**) but using the *maximum* rate of spontaneous events during the same control trials. (**I**) A significant difference emerges in event rates by morphology (n = 23 animals, n = 32, 55, and 8 cells). (**J**) Across animals, Split PCs have higher maximum event rates than Normative PCs (n = 13 animals). (**K**) Animals with a larger difference between local event ratios of split PCs relative to Normative PCs also show a higher event rate in Split PCs (n = 13 animals). (**L**) The maximum rate above the minimum local cell rate is higher in PCs with more local events (n = 23 animals, n = 94 cells). (**M**) PCs with higher local event ratios, particularly Split and Poly, also have elevated event rates (n = 23 animals, n = 95 cells). (**N**) The difference in result between using mean and maximum event rate is partly explained by the fact that Split PCs have wider variability in event rates, with mostly higher maximums and slightly lower minimums, suggesting a larger dynamic range of activity. This is confirmed by computing the gap between minimum and maximum spontaneous event rates during control trials for each cell and comparing Split and Normative PCs (n = 17 animals, n = 13, 16, 6). (**O**) The event rate gap also correlates with the local event ratio (n = 23 animals, n = 95 cells). (**P**) PCs with the minimum spontaneous event rate in each animal (orange) have lower local event rates than the remaining cells (grey, n = 17 animals, n = 17 and 77 cells).

**Fig. S6.**
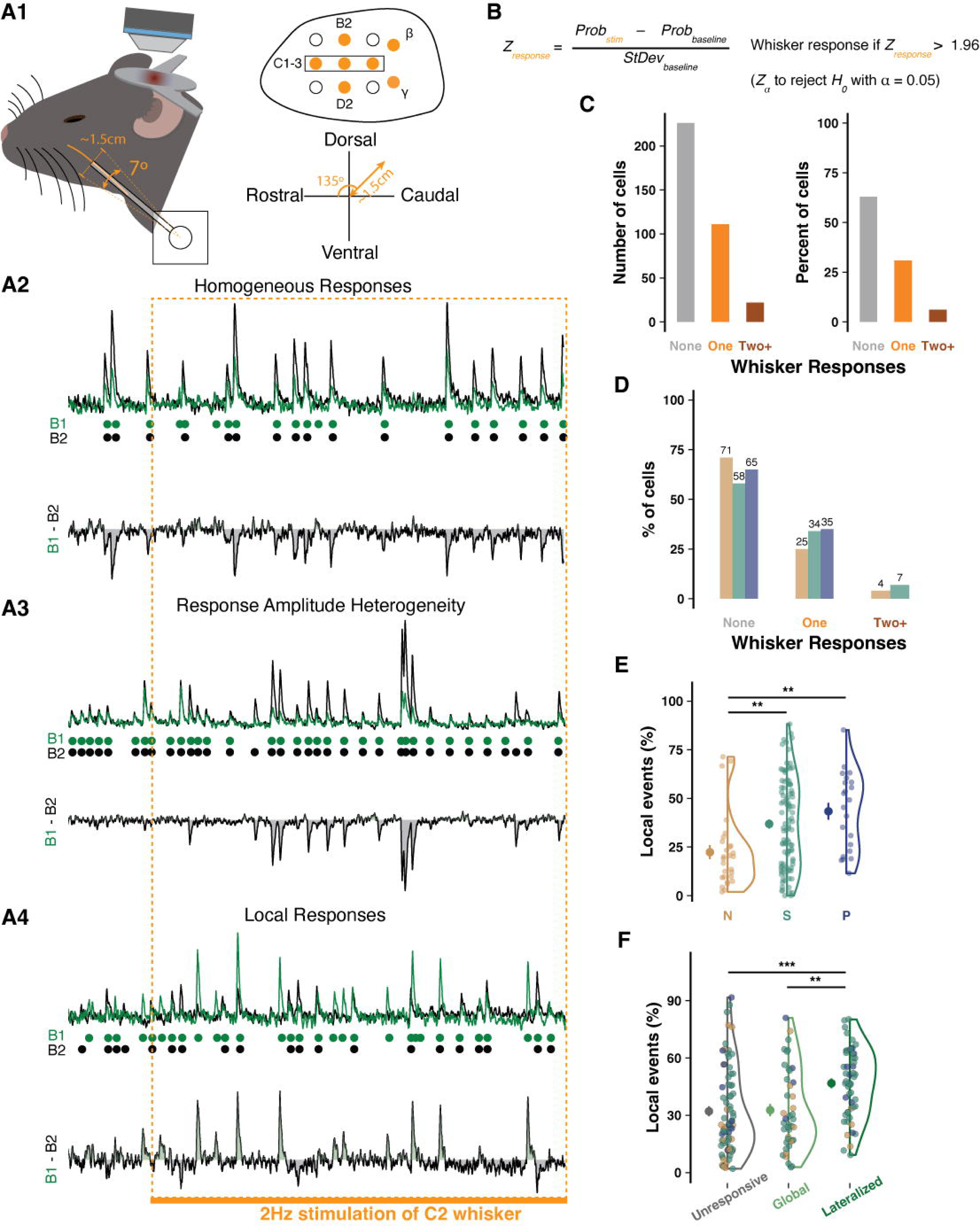
Purkinje cell responses to single whisker stimulation match the literature and branch-specific events match other recording methods. (**A**) Schematic of rotary servo motor stimulation of individual whiskers. Whiskers B2, D2, C1-3, β, and γ were stimulated one at a time for two trials each in a random order in each mouse. The servo motor was programmed via an Arduino synchronized to the imaging time to rotate 7° at 2Hz and was positioned to move each whisker in the dorso-caudal direction (135°). (**A2**) Sample traces from a PC with homogeneous activity and responses during the same trial of stimulation of the C2 whisker as shown in different PCs with either largely global responses but dynamic amplitude scaling (**A3**) or with many local responses in addition to dynamic global response amplitude scaling (**A4**). (**B**) The formula used to determine if events in an ROI constitute a response. A z-scored probability of response is obtained by finding the difference between the probability of an event during baseline vs. response time windows and dividing by the deviation of the probability during baseline periods. If the Z-score exceeded two standard deviations, Z_response_ > 1.96, we could reject H_0_ that there was no response with a confidence of α = 0.05). (**C**) Relative rates of non-response (60%) and response to one (25%) or more whiskers (15%) for cells using combined (averaged) branch signal, the typical approach in the field, as opposed to total responses extracted separately from each branch. These values match the expected rate described by Ju and colleagues (42). (**D**) The relative rates as in (**C**) separated by cell morphology. A slight elevation in response rate can be observed in Split and Poly PCs, even when only assessing the averaged dendritic signal. As shown in Fig. 4D, deconvolving events and responses separately for each primary dendrite in the same cell population produced a higher fidelity representation of whisker responses, which revealed more responsiveness than detected with averaged dendritic signal. Crucially, this elevation of detected responsiveness was especially pronounced in Split and Poly PCs, while Normative PC response rates were largely unchanged. (**E**) Rates of spontaneous, branch-specific events by morphological category during recordings in medial Crus I while mice were under anesthesia (n= 13 animals, n = 28, 99, 22 cells). These rates are nearly identical to rates in awake conditions recorded from lateral Crus I. (**F**) PCs with lateralized responses have more branch-specific events (45%) than PCs with global or no response (both 25%, n = 75, 42, 52 cells).

**Fig. S7.**
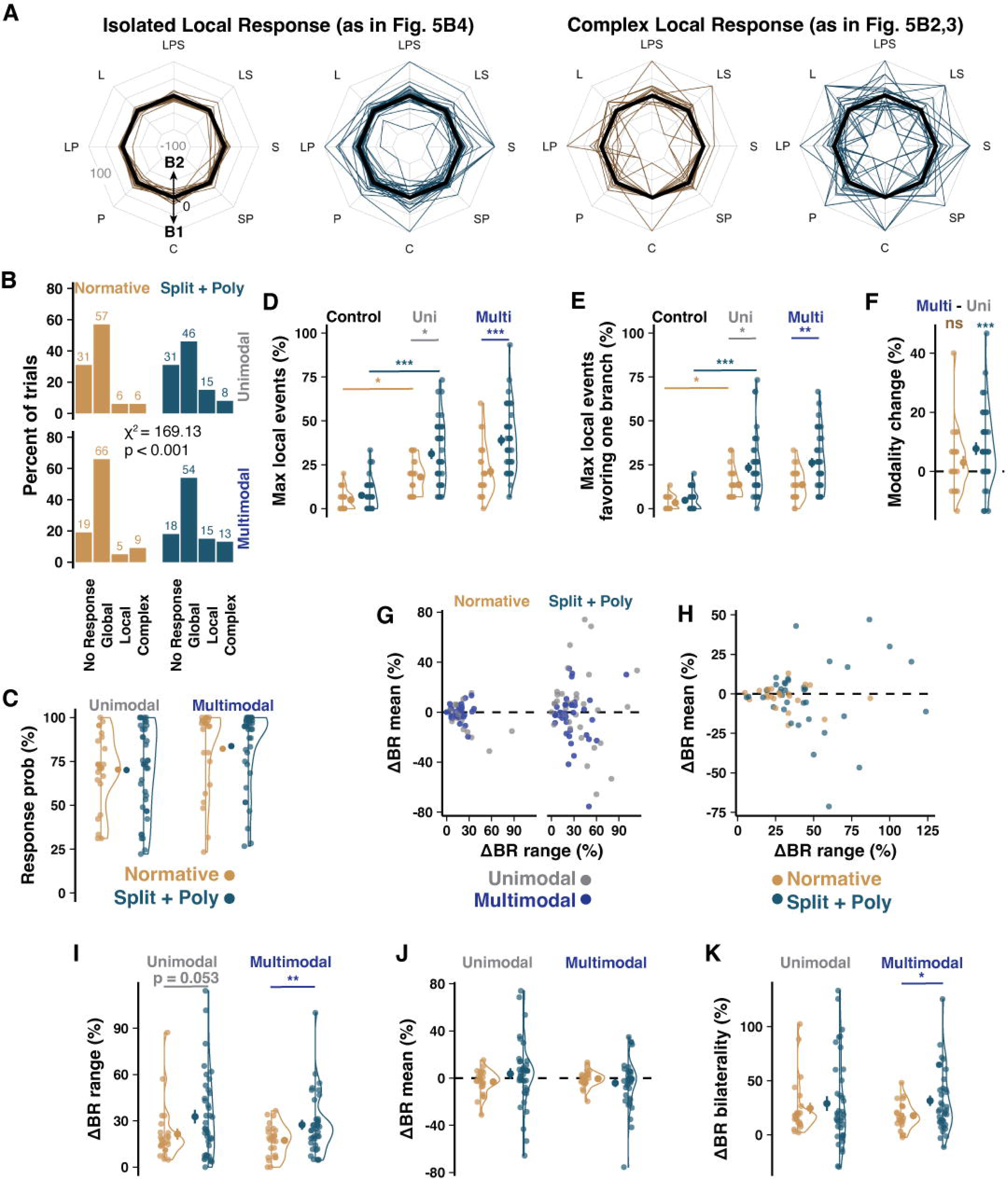
Multisensory CF receptive field representation across Purkinje cell branches. (**A**) Radar plots of non-absolute (directional) local event rates (in isolation, *top*, or as part of a response with global and local components, *bottom*) favoring arbitrarily designated branch 1 or branch 2 of each PC. Middle back line marks no local responses whereas lines deviating toward the outside or inside of the radar indicate high ratios of local events favoring one or the other branch. (**B**) Ratios of uni- and multisensory response type by morphology (n = 2,520 and 3,990 events, from 24 and 38 cells). (**C**) Average response probability (here and below: n = 12 animals, n = 24 Normative and 38 Split+Poly cells). (**D**) As in (Fig. 5C), the highest percentage of local events regardless of branch identity, across stimuli for each cell. (**E**) Obtaining the difference between the number of local responses from each branch for each stimulus – if both branches have local responses to a stimulus this determines how many more local responses one branch had over the other – gives the directionality of the local responses as favoring one branch or the other. Taking the maximum absolute value gives the maximum directional rate. (**F**) Relative difference of maximum local events between uni- and multisensory stimuli reveals no change in Normative PCs but a modest increase of local events in SP PCs during multisensory stimulation. (**G-H**) The relationship between range and directional ΔBR mean grouped by sensory modality category (uni vs. multi-sensory) or morphology (Normative vs. Split+Poly). (**I-J**) As in (Fig. 5F-G), the ΔBranch Response (ΔBR) range and mean, but here separated by uni-vs multisensory stimulus types instead of combined (Student’s t-test). (**K**) As in (Fig. 5H), subtracting the ΔBR mean from the range distinguishes cells with either unilateral or bilateral profile of modality representations. Unilateral cells (high ΔBR range and mean) have one branch that shows branch-specific responses to some but not all modalities, whereas both branches of bilateral cells (high ΔBR range and low ΔBR mean) show branch-specific responses to different modalities.

**Table S1.**
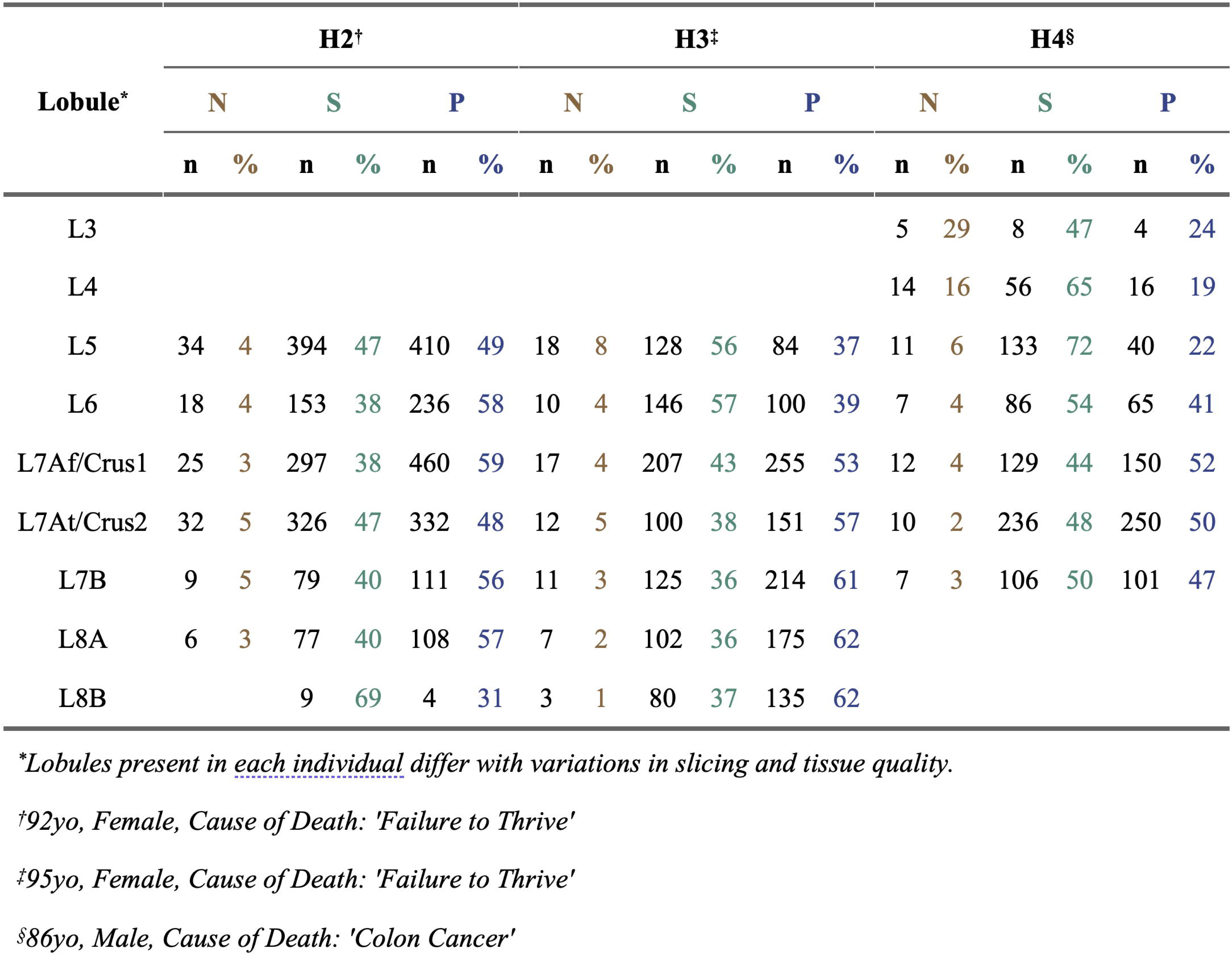
Distribution of PC morphologies in human. Lobule specific numbers and percentages (in color) of each morphological type for each individual in the study. Lobule information is empty if the lobule was not present or could not be assessed in sections from that individual. Related to Figure 1B-C.

**Table S2.** Distribution of PC morphologies in mouse. Lobule specific numbers and percentages (in color) of each morphological type for each mouse in this experiment. Related to Figure 1E-F.

| Comparison | Difference | Lower | Upper | Adjusted p value |
| --- | --- | --- | --- | --- |
| Split:M:A-Normative:M:A | -20.98 | -34.72 | -7.25 | 0.00 |
| Poly:M:A-Normative:M:A | -45.03 | -59.07 | -30.98 | 0.00 |
| Normative:H:A-Normative:M:A | -42.92 | -60.83 | -25.02 | 0.00 |
| Split:H:A-Normative:M:A | 1.79 | -16.11 | 19.70 | 1.00 |
| Poly:H:A-Normative:M:A | -25.77 | -43.67 | -7.86 | 0.00 |
| Normative:M:P-Normative:M:A | -15.18 | -27.71 | -2.64 | 0.01 |
| Split:M:P-Normative:M:A | -17.74 | -30.27 | -5.20 | 0.00 |
| Poly:M:P-Normative:M:A | -33.98 | -46.52 | -21.44 | 0.00 |
| Normative:H:P-Normative:M:A | -52.11 | -65.14 | -39.09 | 0.00 |
| Split:H:P-Normative:M:A | -10.97 | -23.82 | 1.87 | 0.18 |
| Poly:H:P-Normative:M:A | -3.59 | -16.43 | 9.26 | 1.00 |
| Poly:M:A-Split:M:A | -24.04 | -38.08 | -10.00 | 0.00 |
| Normative:H:A-Split:M:A | -21.94 | -39.84 | -4.03 | 0.00 |
| Split:H:A-Split:M:A | 22.78 | 4.87 | 40.68 | 0.00 |
| Poly:H:A-Split:M:A | -4.78 | -22.69 | 13.12 | 1.00 |
| Normative:M:P-Split:M:A | 5.81 | -6.73 | 18.34 | 0.93 |
| Split:M:P-Split:M:A | 3.25 | -9.29 | 15.78 | 1.00 |
| Poly:M:P-Split:M:A | -13.00 | -25.53 | -0.46 | 0.03 |
| Normative:H:P-Split:M:A | -31.13 | -44.16 | -18.10 | 0.00 |
| Split:H:P-Split:M:A | 10.01 | -2.83 | 22.86 | 0.30 |
| Poly:H:P-Split:M:A | 17.40 | 4.55 | 30.24 | 0.00 |
| Normative:H:A-Poly:M:A | 2.11 | -16.04 | 20.25 | 1.00 |
| Split:H:A-Poly:M:A | 46.82 | 28.68 | 64.96 | 0.00 |
| Poly:H:A-Poly:M:A | 19.26 | 1.12 | 37.40 | 0.03 |
| Normative:M:P-Poly:M:A | 29.85 | 16.98 | 42.72 | 0.00 |
| Split:M:P-Poly:M:A | 27.29 | 14.41 | 40.16 | 0.00 |
| Poly:M:P-Poly:M:A | 11.05 | -1.83 | 23.92 | 0.17 |
| Normative:H:P-Poly:M:A | -7.09 | -20.44 | 6.27 | 0.83 |
| Split:H:P-Poly:M:A | 34.05 | 20.88 | 47.23 | 0.00 |
| Poly:H:P-Poly:M:A | 41.44 | 28.26 | 54.61 | 0.00 |
| Split:H:A-Normative:H:A | 44.71 | 23.44 | 65.99 | 0.00 |
| Poly:H:A-Normative:H:A | 17.15 | -4.12 | 38.43 | 0.25 |
| Normative:M:P-Normative:H:A | 27.74 | 10.74 | 44.75 | 0.00 |
| Split:M:P-Normative:H:A | 25.18 | 8.18 | 42.19 | 0.00 |
| Poly:M:P-Normative:H:A | 8.94 | -8.06 | 25.95 | 0.84 |
| Normative:H:P-Normative:H:A | -9.19 | -26.56 | 8.18 | 0.84 |
| Split:H:P-Normative:H:A | 31.95 | 14.71 | 49.18 | 0.00 |
| Poly:H:P-Normative:H:A | 39.33 | 22.10 | 56.57 | 0.00 |
| Poly:H:A-Split:H:A | -27.56 | -48.84 | -6.29 | 0.00 |
| Normative:M:P-Split:H:A | -16.97 | -33.98 | 0.03 | 0.05 |
| Split:M:P-Split:H:A | -19.53 | -36.54 | -2.53 | 0.01 |
| Poly:M:P-Split:H:A | -35.77 | -52.78 | -18.77 | 0.00 |
| Normative:H:P-Split:H:A | -53.91 | -71.28 | -36.54 | 0.00 |
| Split:H:P-Split:H:A | -12.77 | -30.00 | 4.47 | 0.37 |
| Poly:H:P-Split:H:A | -5.38 | -22.62 | 11.85 | 1.00 |
| Normative:M:P-Poly:H:A | 10.59 | -6.41 | 27.60 | 0.64 |
| Split:M:P-Poly:H:A | 8.03 | -8.98 | 25.03 | 0.92 |
| Poly:M:P-Poly:H:A | -8.21 | -25.22 | 8.79 | 0.90 |
| Normative:H:P-Poly:H:A | -26.35 | -43.72 | -8.98 | 0.00 |
| Split:H:P-Poly:H:A | 14.79 | -2.44 | 32.03 | 0.17 |
| Poly:H:P-Poly:H:A | 22.18 | 4.95 | 39.41 | 0.00 |
| Split:M:P-Normative:M:P | -2.56 | -13.77 | 8.65 | 1.00 |
| Poly:M:P-Normative:M:P | -18.80 | -30.02 | -7.59 | 0.00 |
| Normative:H:P-Normative:M:P | -36.94 | -48.70 | -25.18 | 0.00 |
| Split:H:P-Normative:M:P | 4.20 | -7.35 | 15.76 | 0.99 |
| Poly:H:P-Normative:M:P | 11.59 | 0.03 | 23.15 | 0.05 |
| Poly:M:P-Split:M:P | -16.24 | -27.45 | -5.03 | 0.00 |
| Normative:H:P-Split:M:P | -34.38 | -46.14 | -22.62 | 0.00 |
| Split:H:P-Split:M:P | 6.76 | -4.79 | 18.32 | 0.73 |
| Poly:H:P-Split:M:P | 14.15 | 2.59 | 25.71 | 0.00 |
| Normative:H:P-Poly:M:P | -18.13 | -29.89 | -6.37 | 0.00 |
| Split:H:P-Poly:M:P | 23.01 | 11.45 | 34.56 | 0.00 |
| Poly:H:P-Poly:M:P | 30.39 | 18.83 | 41.95 | 0.00 |
| Split:H:P-Normative:H:P | 41.14 | 29.05 | 53.23 | 0.00 |
| Poly:H:P-Normative:H:P | 48.53 | 36.44 | 60.62 | 0.00 |
| Poly:H:P-Split:H:P | 7.39 | -4.51 | 19.28 | 0.65 |

**Table S3.**
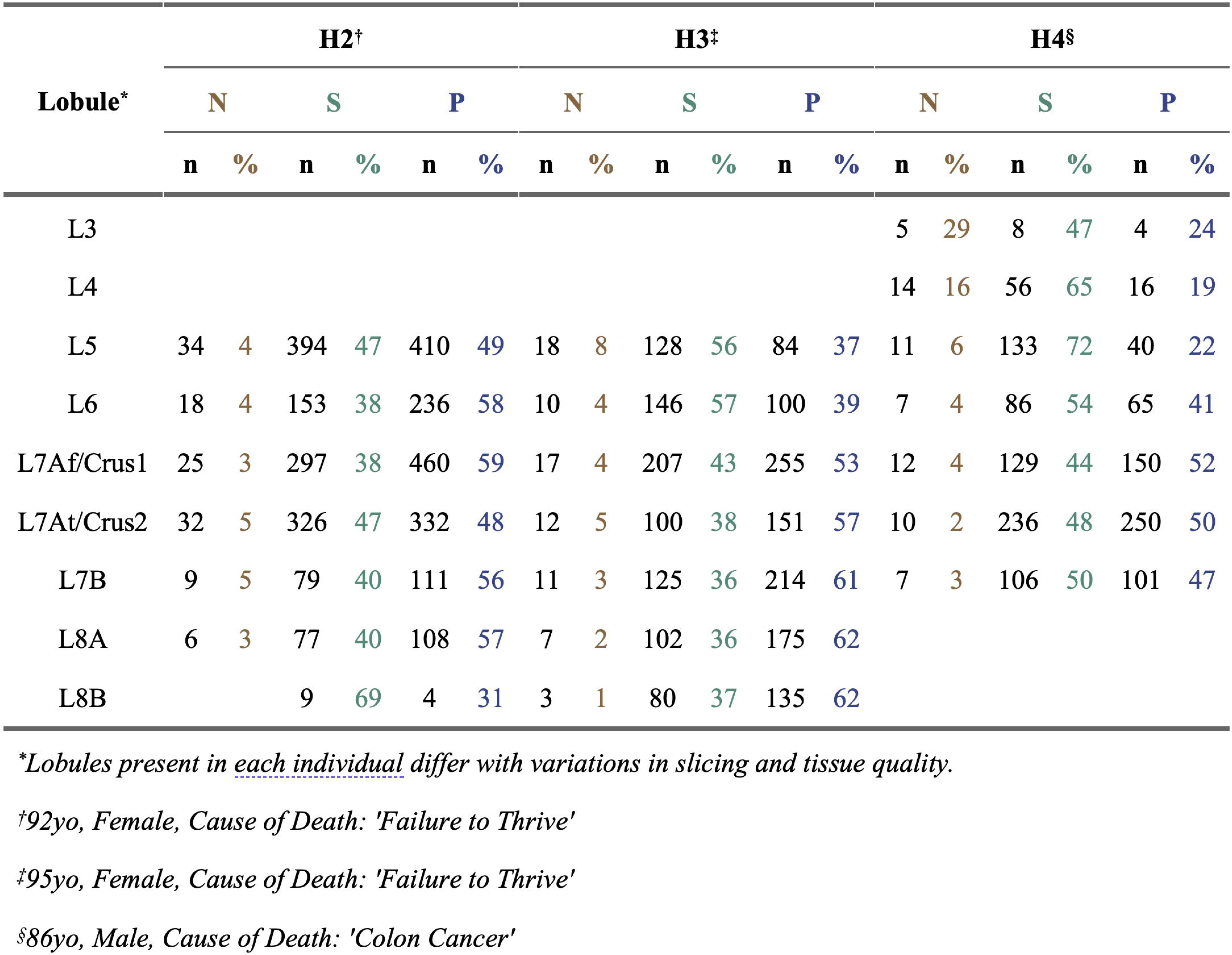
Morphology demographics by cerebellar region and species. Quantification of the results of a three-way ANOVA with TukeyHSD correction for multiple comparisons of morphological demographics by cerebellar region and species. M = Mouse; H = Human; A = Anterior lobules (L2-L5); P = Posterior lobules (L6-L10). Related to Figure 1G and fig. S2A.

**Movie S1. Sample video of spontaneous calcium signal heterogeneity.**

An example of sparse expression of GCaMP6f yielding a lone Purkinje cell in the mouse. Individual branches exhibit global events as well as events that are local to either one branch or the other. Video playback speed is reduced to 0.5x speed.

